# Immune transcriptomic differences in paediatric patients with SARS-CoV-2 compared to other lower respiratory tract infections

**DOI:** 10.1101/2025.11.07.687132

**Authors:** Negusse Tadesse Kitaba, Lesley Workman, Cheryl Cohen, Diana Baralle, Ellen Kong, Maresa Botha, Marina Johnson, David Goldblatt, Mark P Nicol, John W Holloway, Heather J Zar

## Abstract

The clinical severity of SARS-CoV-2 infection in children varies, with asymptomatic or mild illness predominating and a minority developing severe disease. Understanding the immunological responses that underlie severity of disease may guide future development of preventive or therapeutic interventions. This study compared whole blood transcriptomes of healthy children (N=127), children with mild/asymptomatic SARS-CoV-2 infection (N=71) and children hospitalised with severe SARS-COV-2 (N=41), lower respiratory tract illness (LRTI) or LRTI due to Respiratory Syncytial Virus (RSV-LRTI) (N=47) or Pulmonary Tuberculosis (PTB) (N=47). We identified >5000 differentially expressed genes including: *OLFM4, IFI27, CBX7, IGF2BP3, OTOF* for severe SARS-CoV-2; *IFI27, OTOF, SIGLEC1, IFI44L* and *USP18* for RSV-LRTI, and *MMP8, LTF, IGF2BP3, GPR84, CD177, C1QC* and *DEFA4* for PTB, at false discovery rate (FDR) <0.05. Pathway analysis identified enrichment for neutrophil degranulation, interferon gamma signalling, overexpression of ribosomal proteins and depletion of immune response in severe SARS-CoV-2 compared to healthy (SAR-COV-2 uninfected) children. Weighted Gene Co-expression Network Analysis (*WGCNA*) identified 10 correlated gene modules shared between LRTI showing similar underlying response mechanisms. Cellular decomposition analysis identified the depletion of 22 cell types in severe SARS-CoV-2, 16 for RSV-LRTI and 21 for PTB compared to healthy SARS-CoV-2 uninfected control children. We identified 82 genes important for discriminating asymptomatic/mild from severe SARS-CoV-2 including *CBX7, TRAF1, ZNF324* and *CASS4*; 93 healthy from severe SARS-CoV-2 including *RORC, CBX7, NR3C2, MID2* and *ADAMTS2*; 110 genes for RSV-LRTI and 95 for PTB children which can be used for future therapeutic targets.

**Graphical abstract:** 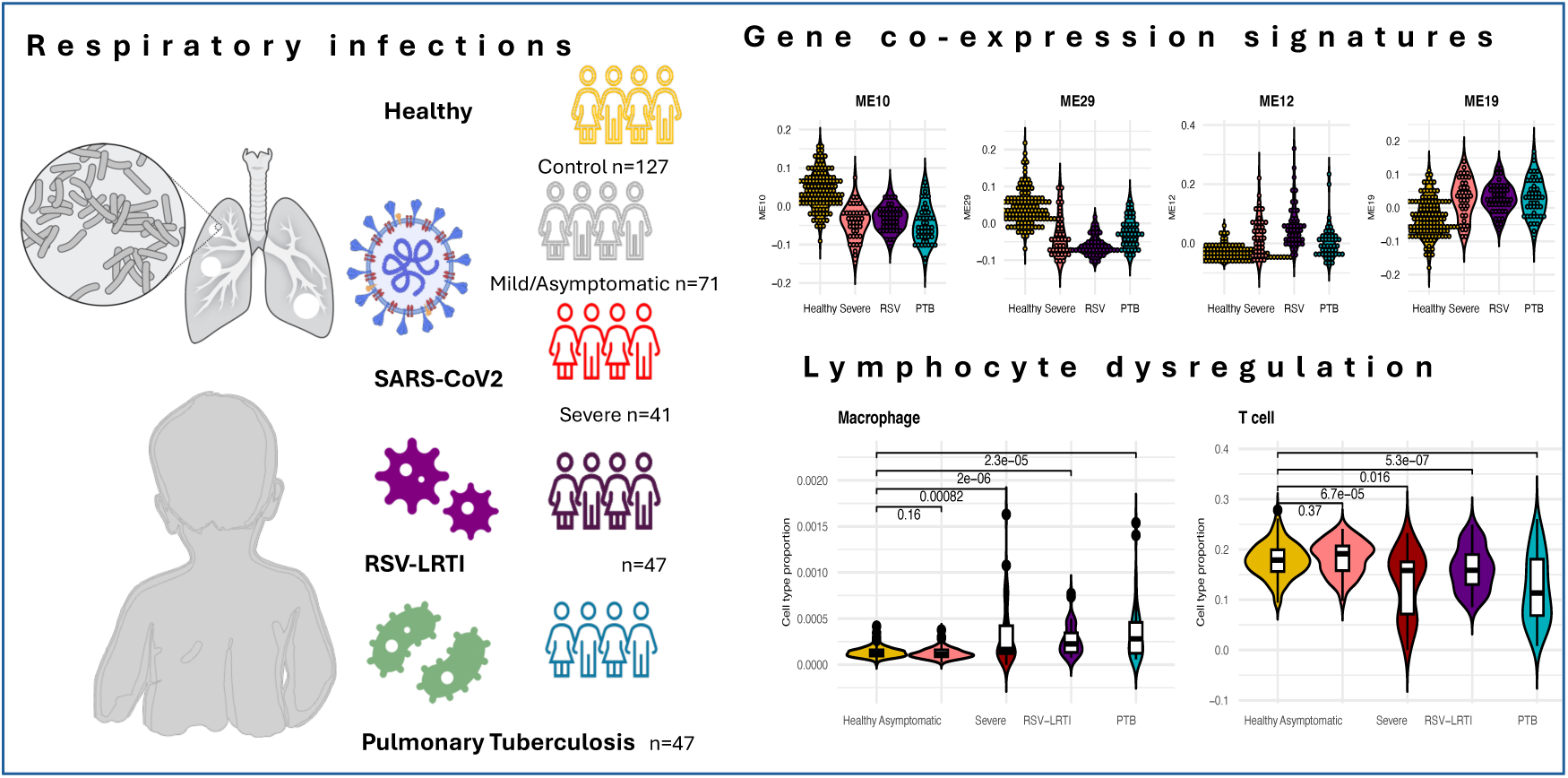

## Introduction

Lower respiratory tract illness (LRTI) is a major cause of hospitalisation and mortality globally in children, with the burden heavily skewed to low-and medium-income countries (LMICs). RSV predominates as a cause of severe LRTI and hospitalisation. Pulmonary tuberculosis (PTB) has also increasingly been recognised as an important cause of acute LRTI in children in countries in which TB is endemic^1^. During the SARS-COV-2 pandemic, SARS-CoV-2 emerged as a cause of LRTI in children.

The clinical manifestation of COVID-19 in children varies widely from mild or asymptomatic illness to severe LRTI^2^, although severe disease is rare. Immunologically, the hallmarks of COVID-19 include dysregulation of type I IFN activity, hyperinflammation, lymphopenia, heterogeneous adaptive immunity, dysregulated myeloid response and lymphocyte impairment^3,4^. COVID-19 severity is also associated with different levels of neutralizing antibodies^5,6^. While the blood transcriptomic response to SARS-CoV-2 infection has been described in adults^7,8,9^, few studies have investigated responses to SARS-CoV-2 in infants and children^10,11^ and little is known about differences in host gene expression between children asymptomatic with SARS-CoV-2 infection and those hospitalized with severe COVID-19 or other LRTI such as Respiratory Syncytial Virus (RSV-LRTI) or pulmonary tuberculosis (PTB)^12,13,14,15^.

A multi-omics approach has previously shown utility in characterising the complexity and severity of Covid-19^16^. Weighted Gene Co-expression Networks Analysis (WGCNA) is a widely implemented approach to identify co-regulated genes and potential hub-genes for druggable targets^17^. The aim of this study was to compare host RNA gene expression in healthy children compared to those with asymptomatic or mild SARS-CoV-2 infection, as well as to those hospitalised with COVID-19, RSV-LRTI or PTB and to utilise WGCNA to identify underlying immune responses associated with disease.

## Methods

This was a prospective study conducted during the SARS-COV-2 pandemic that investigated patterns of whole blood gene expression in HIV-negative children enrolled in a South African birth cohort study, the Drakenstein Child Health study (DCHS), and those hospitalised with SARS-COV-2 (severe COVID-19), RSV-LRTI or PTB.

### Participants

#### Healthy controls or previous SARS-CoV-2 mild or asymptomatic infection

Participants were from the Drakenstein Child Health Study, a prospective population-based birth cohort study of children in a low- and middle-income, peri-urban community outside Cape Town, South Africa^18^. In the DCHS, during the SARS-CoV-2 pandemic, a convenience sample of a subset of children (N=201) was included in intensive surveillance for SARS-CoV-2 infection with blood sampling every 3 months from 15-May-2020 through 15-Sept-2022, with blood and nasopharyngeal swabs collected, irrespective of symptoms.

In addition, continuous surveillance for illness or hospitalisation was undertaken, and blood and nasal sampling repeated at any intercurrent illness. Serum samples were stored and batched for measurement of IgG to Spike antigen (CoV-2-S-IgG) by ELISA as previously described^19^. In the current study, samples from children during wave 1 were used; subjects seronegative for SARS-CoV-2 were defined as healthy controls, and those seropositive for SARS-CoV-2 were considered mild/asymptomatic infection as no child reported symptomatic illness or was hospitalised.

### Children with LRTI

#### COVID-19 or RSV-LRTI

Children with acute LRTI hospitalised at Red Cross Childrens Hospital were identified through the National Syndromic Surveillance for pneumonia in South Africa programme (PSP) at Red Cross War Memorial Children’s Hospital, in Cape Town, South Africa. Sequential children hospitalised with LRTI were enrolled and a nasal swab for PCR detection of SARS-CoV2, RSV and other pathogens was taken for testing at National Institute of Communicable Disease as previously described^20^. Children who were positive for SARS-CoV-2 and negative for other pathogens were considered to have severe COVID-19 (N=41); those positive for RSV were included as RSV-LRTI (N=51).

#### PTB

Children enrolled in a TB diagnostic study (N=47) at Red Cross Children’s Hospital, microbiologically confirmed (by mycobacterial liquid culture or Xpert MTB/RIF) and negative for SARS-CoV-2 and RSV, were included in this study. Serum and PAXgene samples were collected at the time of illness (Severe COVID-19, RSV-LRTI, PTB) were used for this study^20^. Whole blood PAXgene samples were stored at -80°C, randomized prior to shipment, with RNA extraction and sequencing undertaken at the Genomics Shared Resource (GSR), Roswell Park Comprehensive Cancer Centre, Buffalo NY, USA.

### Sequencing and processing RNAseq data

Raw reads were processed with the bcbio-nextgen pipeline. Reads quality were assessed using FastQC^21^ and MultiQC^22^. Sequencing reads were aligned to the human transcriptome reference using STAR^23^. Quantification of gene expression was carried out using Salmon^24^ with default settings. Read counts were normalized using CPM (counts per million) from edgeR^25^ with the TMM (Trimmed Mean of the M-values) method which accounts for both sequencing depth and gene length^26^. Sample outliers were detected using Robust Principal Component Analysis (rPCA) with PcaHubert and PcaGrid functions^27^; samples detected by both methods were excluded from downstream analysis.

### Identification of differentially expressed genes (DEGs)

Amongst 198 children in DCHS, 64% were seronegative (N=127) and regarded as healthy controls. Those were compared to hospitalised children with COVID-19 (N=41), RSV-LRT (N=47) or PTB (N=47). SARS-CoV-2 seropositive during wave 1 (N=71), who did not report any respiratory symptoms or hospitalization over this period, were regarded as having had mild or asymptomatic infection.

The R-package limma^28^ was used to identify differentially-expressed genes adjusting for children’s sex and age. Multiple testing correction was performed using the Benjamini-Hochberg (BH) procedure for False Discovery Rate (FDR) < 0.05. The biological function of gene lists were identified via gene set and pathway enrichment analyses using toppGene^29^.

### Weighted Gene Co-expression Network Analysis (*WGCNA*)

Signed weighted gene co-expression network analyses were conducted using WGCNA^30^. The gene module/clusters represent genes with highly correlated expression patterns, where the first principal component of the gene expression profile (Eigengene) is used to summarise the overall expression of each module. The module eigengenes identified by WGCNA were correlated with Severe COVID-19, PTB and RSV-LRTI. The module associations were visualised as a correlation barplot using the lares R package^31^. Protein-Protein Interaction (PPI) network were identified with GeneMANIA^32^ and network properties for hub genes were analysed and visualized using Cytoscape^33^. Significantly associated modules were further characterized for functional enrichment using toppGene^29^. Non-redundant biological process terms were generated and visualized using rrvgo package^34^.

### Cell type proportion estimation

Cell type proportion differences between groups were estimated and assessed using xCell 2.0^35^ using the Immune Compendium^36^ and immunoprofiling^37^ reference datasets. The t-test was used to determine the difference between groups (asymptomatic vs hospitalized SARS-CoV-2, control vs RSV-LRTI and control vs PTB).

### Severity predictors

Gene biomarkers to predict SARS-CoV-2 severity, RSV-LRTI or PTB were selected using the Boruta^38^ R package^39^ with default settings.

### Gene and target drug look-up

In order to identify the druggability of differentially expressed genes, the look-up target score generated by DrugnomeAI^40^ was utilised (accessed on 19 March 2025). All statistical analyses were conducted in R version 4.5.1.

## Results

### Participant characteristics

This analysis includes 333 children: 71 with previous mild/asymptomatic SARS-CoV-2, 127 seronegative, healthy, and 135 children hospitalised with LRTI (41 with SARS-COV-2, 47 with RSV-LRTI and 47 with PTB or pulmonary TB). The characteristics of each group are shown in Table 1. As there was a significant age difference between DCHS children and those with LRTI, age was included as a covariate in regression analyses.

**Table 1.**
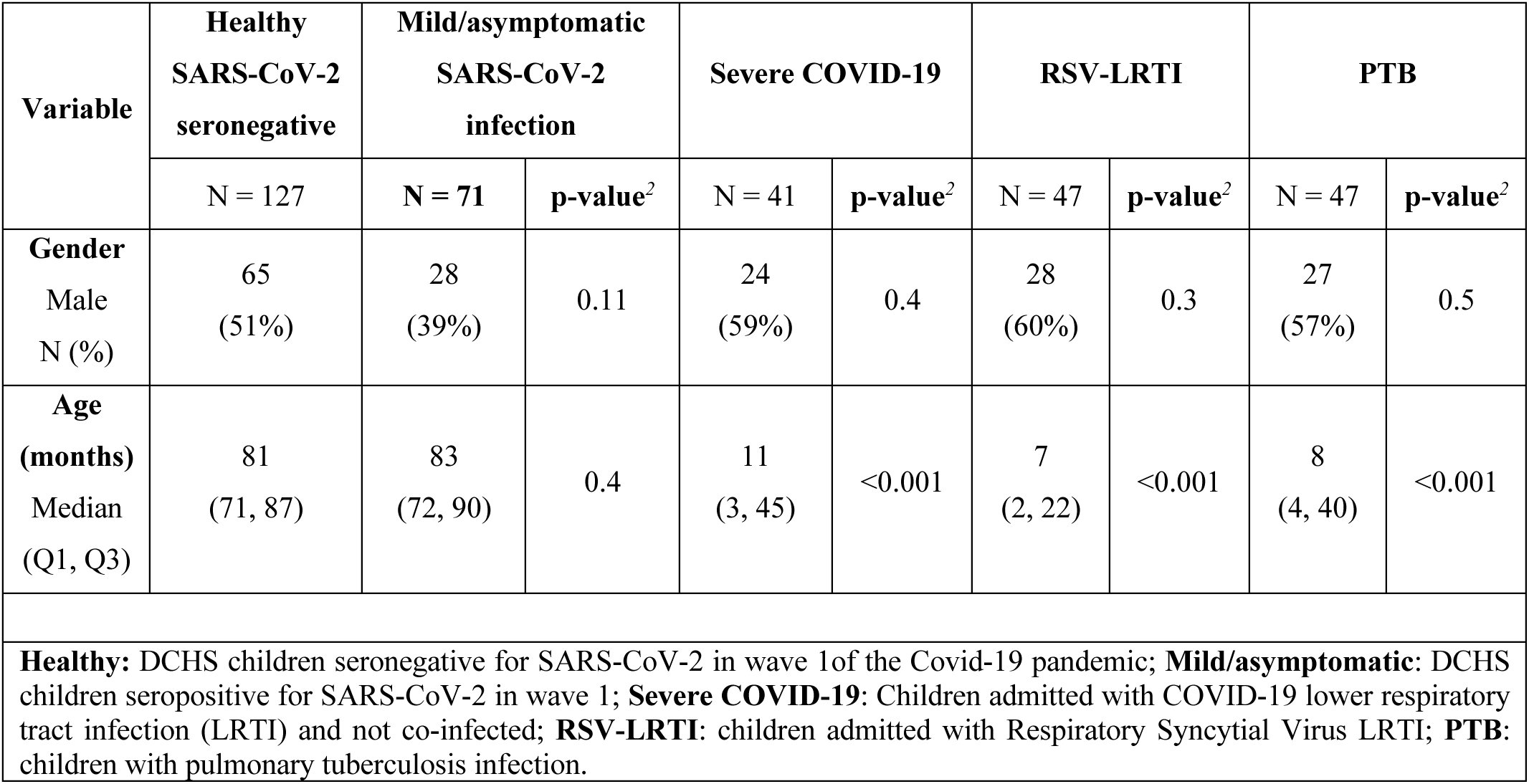
Comparison of participants’ characteristics for healthy controls and children with respiratory tract infections.

### Differential gene expression analysis

To identify differentially expressed genes and enriched GO terms in children with LRTI, seronegative DCHS participants from wave 1 (healthy controls) were compared to each LRTI group separately (COVID-19, RSV-LRTI, PTB). The summary statistics and gene lists for TWAS at FDR <0.05 are provided in Supplementary Table S1. The biological gene ontology enrichment is also provided in Supplementary Table S2.

### COVID-19 disease

The transcriptional response in healthy controls was compared to hospitalised children with COVID-19. There were 118 up-regulated and 160 down-regulated differentially expressed genes (DEGs) between healthy control and severe SARS-CoV-2 cases (FDR < 0.05 and log2 fold change >1), as shown in Figure 1A. Top DEGs included: *IFI27, MMP8, OLFM4, CEACAM8, LTF, IGF2BP3, DEFA4, ADAMTS2* and *CBX7*. Pathways identified as enriched include regulation of immune system and lymphocyte activation (Figure 2A).

**Figure 1.**
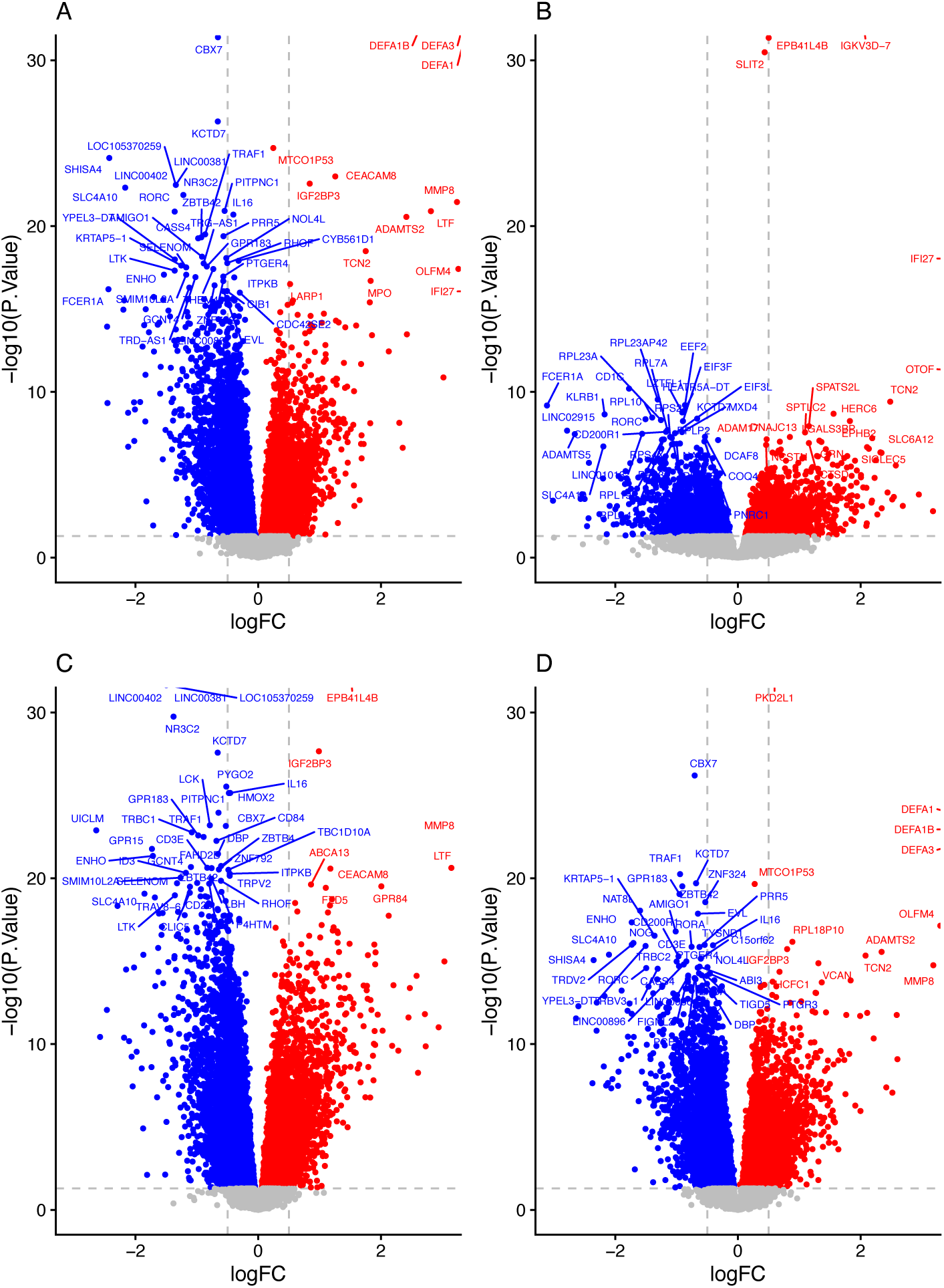
TWAS Volcano plot. A) Severe COVID-19, B) RSV-LRTI, C) PTB D) COVID-19 Severity. Red - upregulated, Blue - down regulated (P value <0.05, log2 Fold Change > 0).

**Figure 2.**
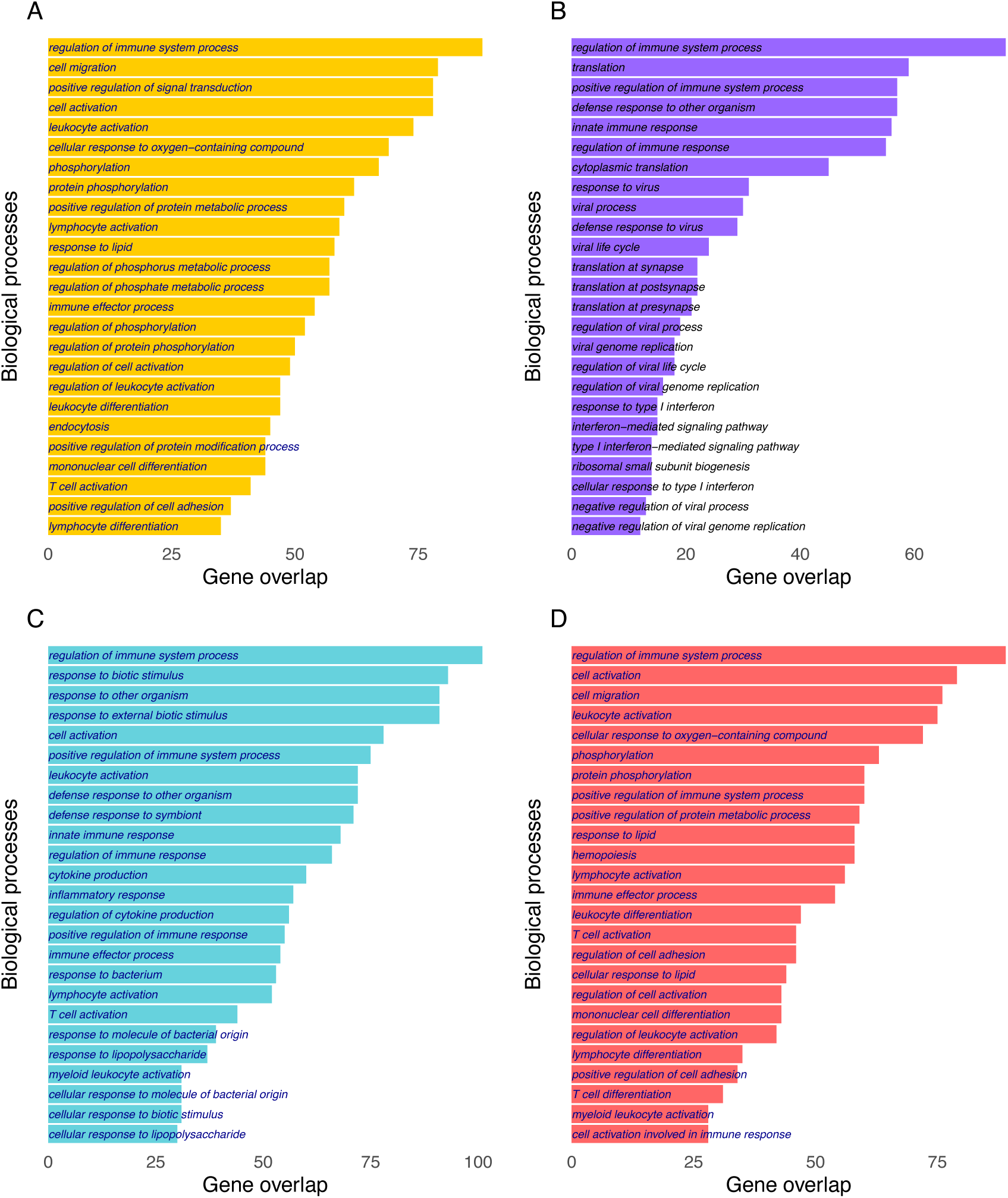
Gene ontology term for biological process for top 500 genes. A) Severe COVID-19 B) RSV-LRTI C) PTB D) COVID-19 severity.

### RSV-LRTI

DGE analysis identified 210 upregulated and 195 downregulated genes at FDR < 0.05 and log2 fold change >1 and differentially expressed between healthy controls and children hospitalized with RSV-LRTI; top DEGs included *IFI27, OTOF, SIGLEC1, IFI44L, USP18, TCN2, CD177, HERC6, C1QC* and *EPHB2*. For all summary statistics see RSV-LRTI in Table S1 and the volcano plot shown in Figure 1B. Pathways significantly enriched included regulation of immune system translation, interferon mediated signalling, viral life cycle and viral processing (Figure 2B).

### Pulmonary Tuberculosis

Children with PTB had identified 203 upregulated and 1843 downregulated genes differentially expressed genes (FDR < 0.05 and log2 fold change >1) compared to healthy controls. Top genes identified include *MMP8, LTF, IGF2BP3, GPR84, CD177, C1QC, DEFA4* and *OLFM4* (see Figure 1 and Supplementary table S1). The pathways identified as enriched include defence response to bacteria, and innate immune response (see PTB in Figure 2C).

### Severity of SARS-CoV-2 infection

In order to determine transcriptional responses that distinguish mild/asymptomatic SARS-CoV-2 infection from severe COVID-19, children hospitalised with COVID-19 were compared with seropositive DCHS children. We identified 163 upregulated and 183 down downregulated genes at FDR < 0.05 and log2 fold change >1 see supplementary table S1. The pathways identified as enriched include regulation of immune system, hemopoieses and lymphocyte activation (see Figure 2D and supplementary table S2).

#### Weighted Gene Co-expression Networks Analysis of LRTI

The WGCNA analysis identified 46 significant modules including 22 with severe COVID-19, 22 with RSV-LRTI and 20 with PTB when compared with healthy controls (p<0.05). Modules 10, 29, 22, 28 and 15 were downregulated and modules 32, 7, 19, 26 and 12 upregulated across LRTIs. The distribution of Eigengenes vs LRTI is shown in Figure 3A and Supplementary Table S3. The distribution of the relationship between the modules is represented as a dendrogram (Supplementary Figure 1) and genes per module are shown in Supplementary Table S3. The correlation of modules with COVID-19, RSV-LRTI and PTB are shown in Figure 3B. Thirty modules showing correlation across LRTIs (r >0.25) were identified, of which 10 modules were correlated with all LRTI, 6 were in common between COVID-19 and RSV-LRTI, and 6 between COVID-19 and PTB (see Table 2 and Supplementary Fig 3). There were 4 modules specific to RSV-LRTI and 2 were specific to PTB.

**Figure 3.**
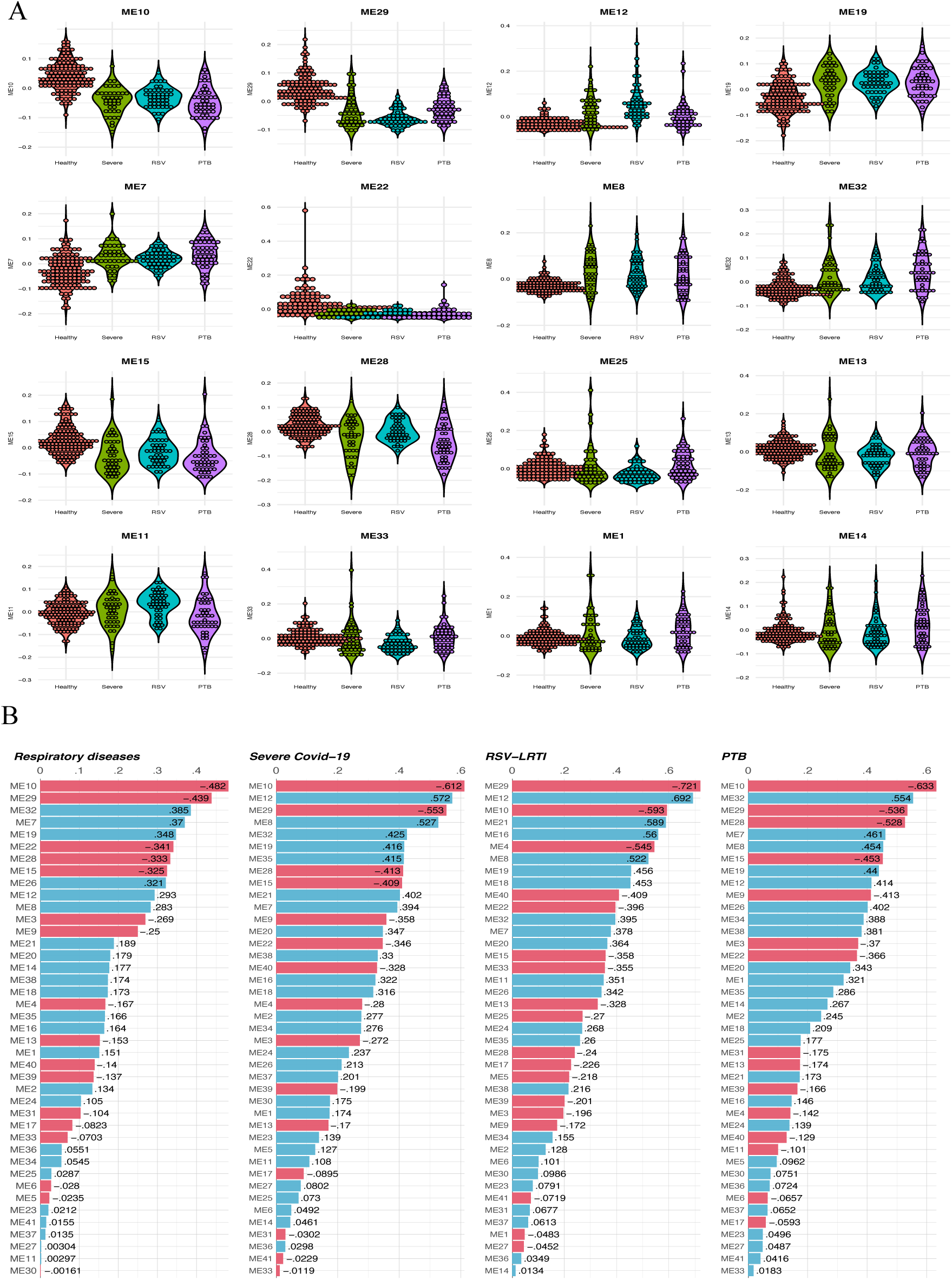
WGCNA analysis for respiratory infections and module correlation. A) Distribution of significant modules per respiratory infection (x-axis module eigengene vs y-axis respiratory infection), B) Correlation of Eigengene with respiratory infection.

**Table 2.**
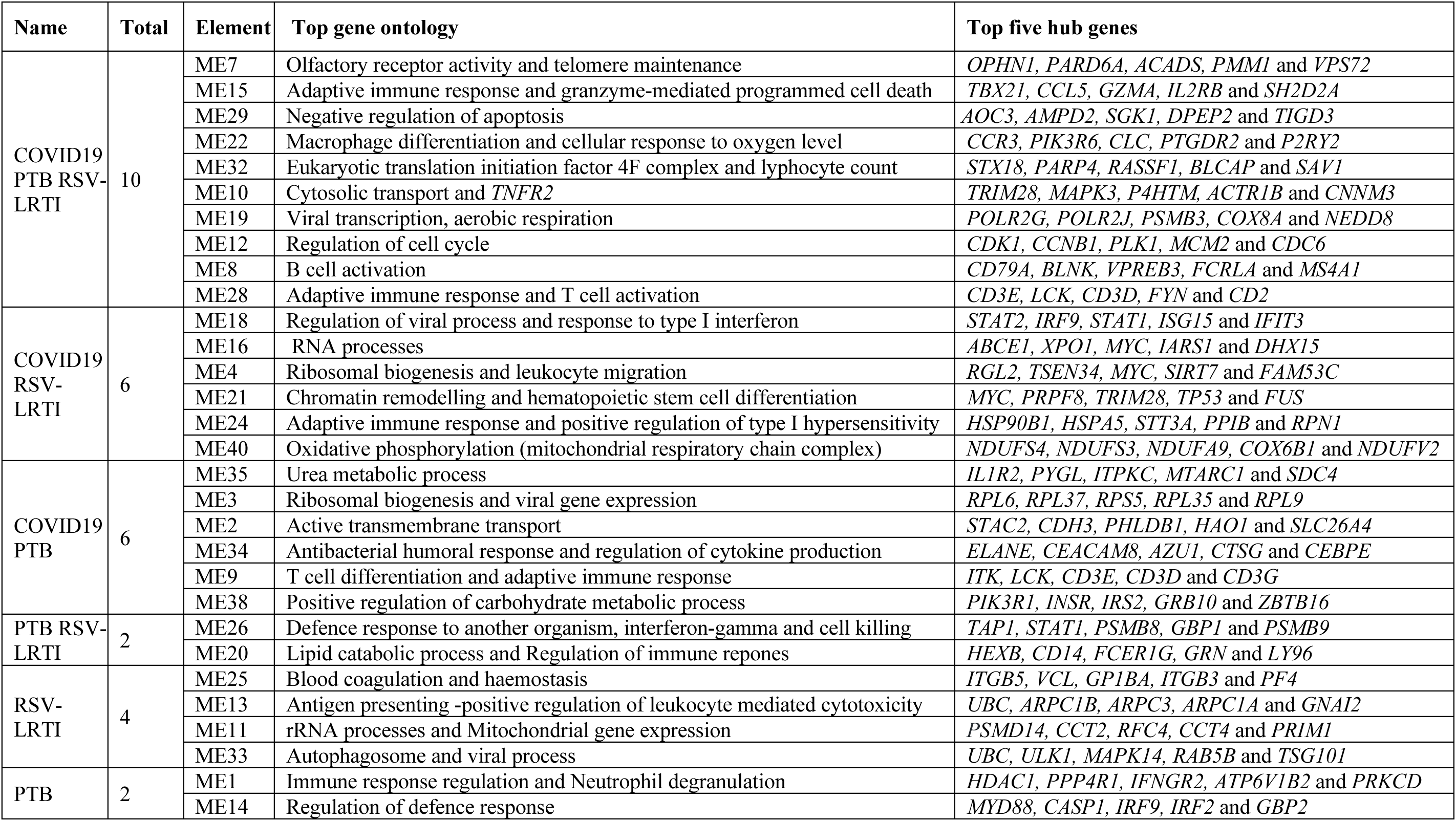
WGCNA modules correlated with respiratory infections at Pearson correlation r >0.25 and p< 0.05.

The gene list in each module was used to generate a network using GeneMANIA with 10 additional interactors for biological processes in Cytoscape. Network analyses were conducted to characterise the network properties including identifying hub genes based on degree of connectivity. The top five hub genes for modules are shown in Table 2. The network connectivity degree distribution for each module is provided in Supplementary Table S4. The Cytoscape session is also provided as Supplementary file 1.

The GO terms enrichment for modules which showed Pearson correlation of r > 0.25 with specific LRTI is in Supplementary Table S5. Further, redundant gene ontology was removed based on similarity matrix of GO terms using rrvgo R package. For biological process visualisation for all other modules see Supplementary Figure 4.

### Cell population differences associated with LRTI

To determine cell type composition differences in peripheral blood between healthy controls and hospitalised subjects with LRTI due to different pathogens, blood cell type proportions were estimated with xCell2 2.0 generated with the ImmuneCompendium.xCell2Ref reference panel. Significant cell type composition differences were identified between healthy controls and hospitalised LRTI groups: 23 for severe COVID-19, 16 for RSV-LRTI and 21 for PTB (p <0.05) (see Fig 4). To determine cell type composition difference between LRTIs we conducted t-tests as shown in Figure 5B. There was no difference in cell composition between healthy controls and those with mild/asymptomatic COVID-19. When the different hospitalised LRTI groups were compared with each other, several differences in cell composition were observed (p< 0.05). These included T cells (lower in severe COVID-19 vs PTB), non-classical monocytes (severe COVID-19 vs RSV-LRTI and RSV-LRTI and PTB) and myeloid cells (RSV-LRTI vs PTB). PTB also showed depletion of central memory CD8+ T Cells and overexpression of granulocytes compared to RSV-LRTI (see Supplementary Fig 5).

**Figure 4.**
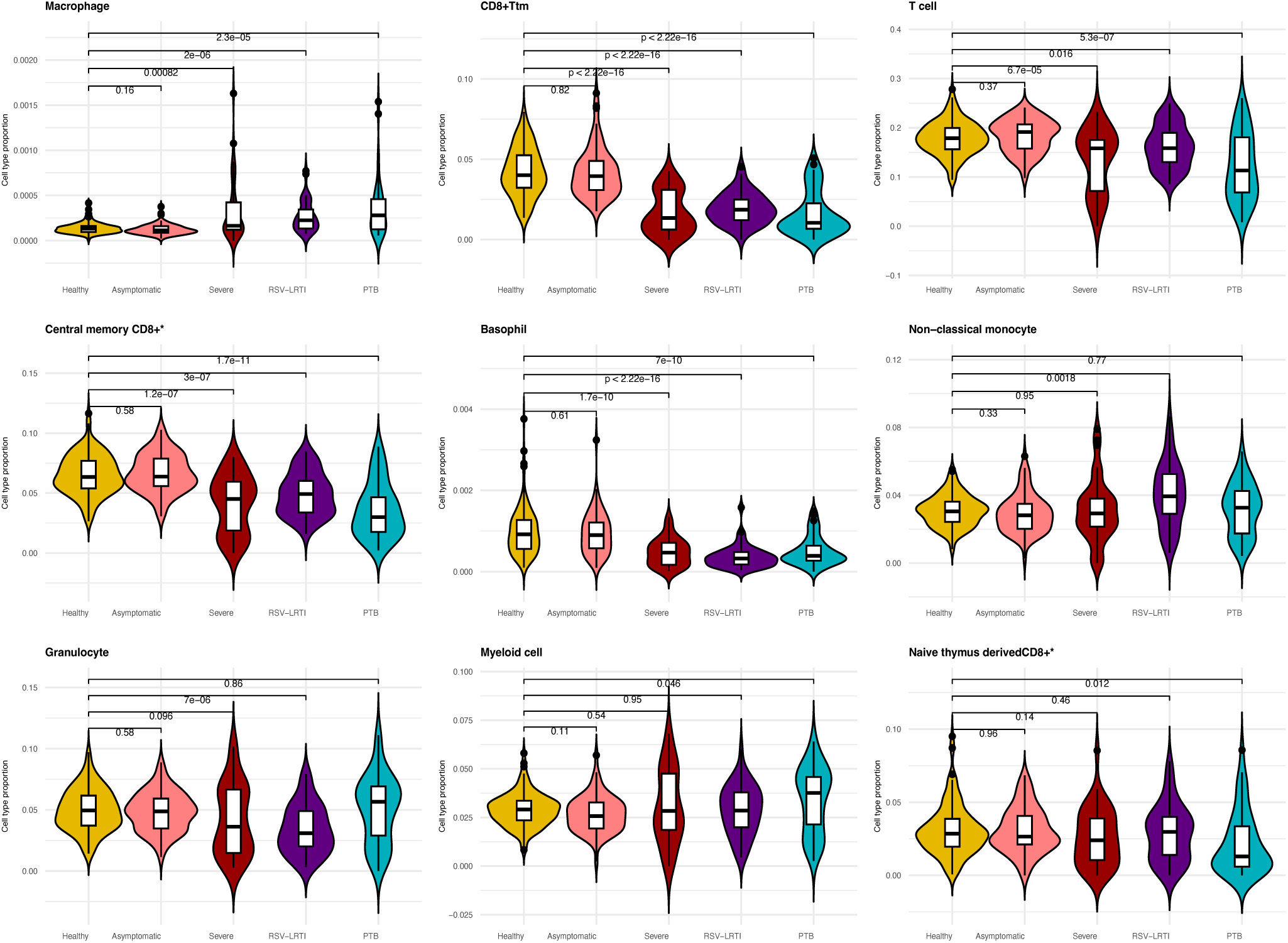
Differences in the proportions of immune cells in respiratory infections comparing healthy controls vs different LRTI groups. It shows the comparison of the five top cell types for LRTI and cell types uniquely different for RSV-LRTI and PTB.

**Figure 5.**
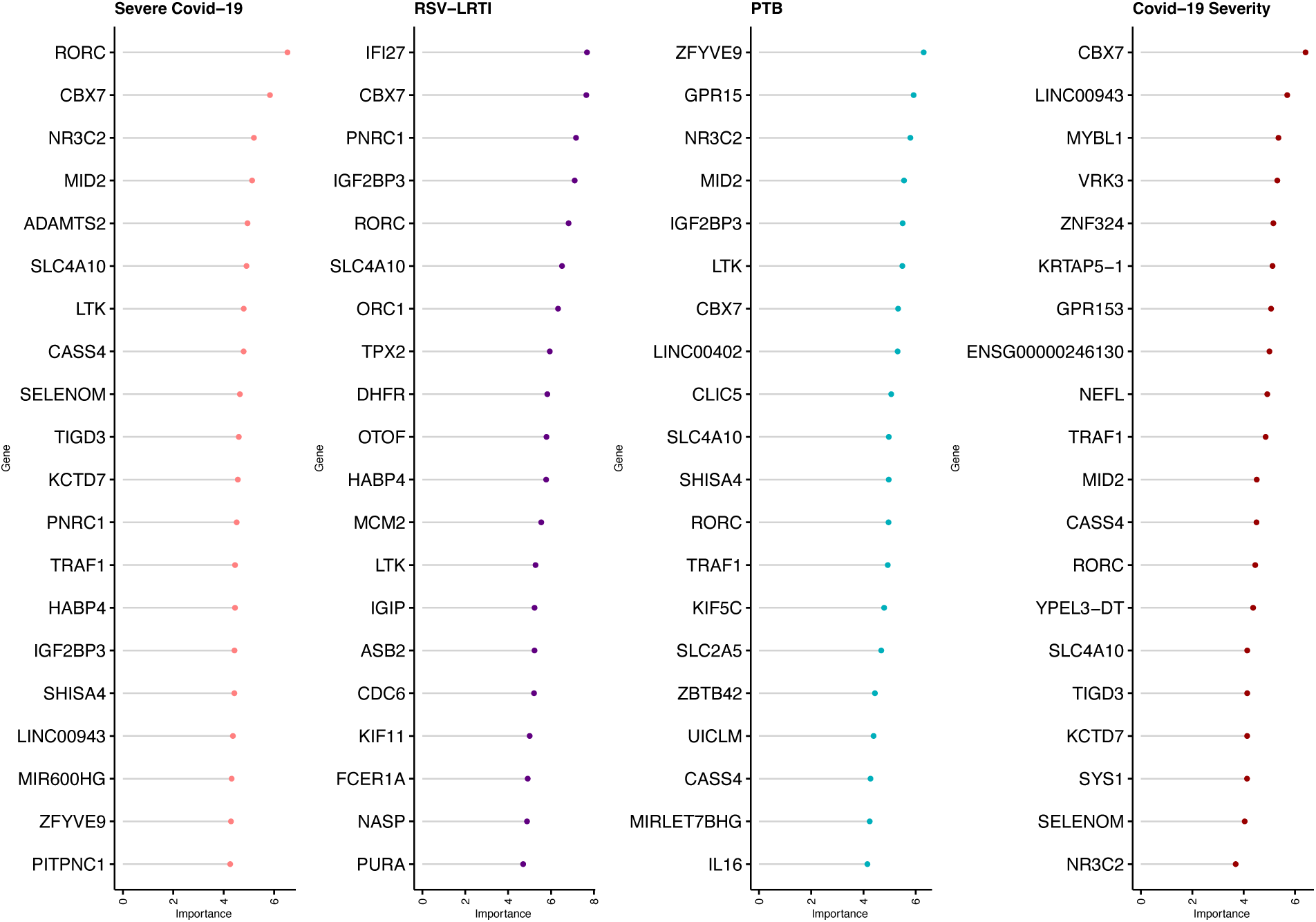
Top 20 gene severity predictors of LRTI for different etiologies based on mean importance. See shared predictors in Supplementary Fig 7.

Seven cell types showed differences with healthy controls across all LRTIs including: Macrophages, transitional memory CD8+ T cells (CD8+ Ttm, T cells, Central memory CD8+ alpha-beta T cells, basophils, myeloid cells and naive thymus-derived CD8+ alpha-beta T cells. Neutrophils and class switched memory B cells showed significant changes for RSV-LRTI and PTB compared to the healthy controls but not for severe COVID-19. Disease specific unique cell type proportion changes were identified for severe COVID-19 as shown in Figure 5. The details are provided in Supplementary Table S6 and Supplementary Fig 6.

### Predictors of severe LRTI

To identify genes that represent biomarkers for each hospitalised LRTI, the normalized counts of the top 1000 significantly differentially expressed genes with respect to healthy controls were used and machine learning algorithms applied to identify the most informative genes. Ninety-three genes were identified as biomarkers for severe (hospitalised) COVID-19, 110 for RSV-LRTI and 95 for PTB as shown in Figure 5 and Supplementary Table S7.

Some genes were able to discriminate specific LRTIs from healthy controls including Severe COVID-19 (23), RSV-LRTI (74), PTB (37) and asymptomatic COVID-19 from severe COVID-19 (COVID-19 severity) (N=25) as shown in the Supplementary Figure 8. There were 10 genes that discriminated healthy controls from any LRTI including *IL16, LTK, IGIP, IGF2BP3, CBX7, KCTD7, FCER1A, TRAF1, RORC* and *SLC4A10*. See details of shared predictor genes amongst the LRTI groups in Supplementary Table S7.

### Drug target lookup for genes associated with LRTI

To determine potentially therapeutic targets from DEGs associated with each LRTI, a look-up was undertaken for overlap with known druggability score generated by drugnomeAI. We identified 689, 159 and 849 genes for COVID-19, RSV-LRTI and PTB respectively with Tclin (approved drug targets) drugnomeAI score >90, as shown in Supplementary Table S8. For availability of drug and new therapeutic options we examined our predictors for availability of drugs as shown in Supplementary Table S9.

## Discussion

The transcriptional landscape of peripheral blood in response to viral and bacterial infections exhibits age-dependent variation, with implications for disease severity and immune regulation. In adults, SARS-CoV-2 infection has been studied extensively and elicits a robust transcriptional response characterized by upregulation of neutrophil activation markers, inflammatory cytokines, and interferon-stimulated genes (ISGs), alongside suppression of adaptive immune pathways and lymphocyte-associated transcripts ^41,42^. Children infected with SARS-CoV-2 typically exhibit mild or asymptomatic disease, with transcriptomic profiles showing restrained inflammatory responses and lower expression of viral entry receptors such as *ACE2* and *TMPRSS2*^43^. However studies of the transcriptional responses to SARS-CoV-2 infection in children are extremely limited, focussing mainly on adolescents^44^. In this study, for the first time, we report genome-wide assessment of transcriptional responses of children hospitalized with one of three LRTIs (COVID-19, RSV-LRTI, PTB), compared to healthy children in a birth cohort from a low- and middle-income African setting. We identify 4500 genes related to hospitalized COVID-19 and known signature genes for RSV-LRTI and for PTB. Unique and shared pathways and gene modules were characterised between LRTIs, along with unique signatures for each of the LRTIs.

COVID-19 related genes were enriched for immune system, neutrophil degranulation and interferon gamma signalling as previously reported in other studies in adults. Neutrophil degranulation has been previously correlated with COVID severity^45^ and excess neutrophil degranulation is associated with tissue damage^46^. The top upregulated genes included known genes responsible for immune responses, such as Interferon alpha-inducible protein 27 (*IFI27*) which is known to be an early predictor for COVID-19 outcome^47^. Many studies have shown reduced ribosomal protein expression and immune suppression associated with persistence of COVID-19 infection^48^. Massoni *et al*^3^ have discussed immune dysregulation and exhaustion as a hallmark of COVID-19 where adaptive immune responses are highly heterogeneous. Thus, at an early phase of infection, type I IFN activity as an anti-viral response is important in the development of both adaptive and innate immunity.

RSV-LRTI upregulated genes include *OTOF*^49^, *SIGLEC1*^50,51^, *USP18*^52^ and *ISG15* ^53^. These genes were enriched for pathways including response to other organisms, regulation of viral life cycle, translational and interferon gamma signalling.

For PTB, we identified genes including *MMP8* and *MMP9* which are known to be associated with TB disease, by degradation of extracellular matrices^54,55,56^. *DEFA1, DEFA1B, DEFA3* and *DEFA4* are a known cluster of genes in the PTB defence response pathway. The expression of *LTF* is also known to be an important biomarker for PTB disease^57,58^. PTB specific markers such as *NCR3, CR2 CD28, IL10RA* and *GPR183* are functionally related to immune response, where *NCR3* stimulates NK cytotoxicity and *CR2* is involved in lymphocyte activation. These findings may contribute to understanding host responses in children in PTB and to strengthening diagnostic possibilities.

Using *WGCNA* co-expression analysis, we identified four RSV-LRTI specific modules: ME11 (translation and aerobic respiration), ME13 (antigen processing and T-cell mediated cytotoxicity), ME25 (coagulation and positive regulation of leukocyte) and ME33 (autophagy, viral processing and negative regulation of ferroptosis). A further two modules were specific to PTB: ME1 (immune response regulating signalling pathway and leukocyte differentiation) and ME14 (regulation of immune and defence response and cytokine production) (see Supplementary Fig 4). While no modules were identified as specific to COVID-19, 22 modules were shared between COVID-19 and one or more LRTI, reflecting the seriousness of SARS-CoV-2 infection.

Ten shared modules were identified across all LRTIs (Table 2) including module 10, which is associated with endosomes^59^ and contains the hub gene *TNFR2*, known to be linked with immune dysregulation in severe COVID-19^60^. In addition, module 10 contains many key hub genes known to be associated with COVID-19 severity including *TRIM28* (265 degree), *P4HTM* (245), *ACTR1B* (244), *CNNM3* (243), and *VPS51* (238). *TRIM28* is known to regulate SARS-CoV-2 entry by targeting *ACE2*^61^, suppressing antiviral immunity^62^ and is linked with COVID-19 severity^63^. *P4HTM* is known to play a role in adaptation to hypoxia and energy response and is linked with hypoventilation^64^.

Other shared modules include: Module 22 the hub gene *CCR3* (C-C motif chemokine receptor 3) regulates cell migration and inflammatory responses by acting as a receptor for various CC chemokines such as eotaxin, and is a susceptibility gene for severe COVID-19 ^65^. Module 28 was related to adaptive immune response and T-cell activation; with hub genes including *CD3E* involved in T-cell signalling to detect and clear pathogens. Module 7 was enriched for sensory perception such as olfactory dysfunction, a known symptom in COVID-19 ^66^. Module 15 was related to T-cell differentiation and adaptive immunity where hub gene *TBX21* is a transcription factor that modulates innate immunity by regulating the expression of *TLR2*^67^. *GZMA* and *GZMB* play a role in immune response during respiratory infection^68^. *IFNG* is involved in clearing viral infection^69^.

A further six modules were shared between COVID-19 and PTB including module 34 which was enriched for antimicrobial humoral responses (*DEFA1, DEFA3, RNASE3, BPI, PGLYRP1, CAMP, AZU1, ELANE* and *LTF*) and neutrophil degranulation^70^ in the Reactome database (*DEFA1, ORM1, ORM2, RNASE3, ATP8B4, STBD1, BPI, PGLYRP1, TCN1, MS4A3, ABCA13, CLEC5A, CAMP, AZU1, CPNE3, CEACAM8, ELANE, CEACAM6, CRISP3, LTF, PLD1, MMP8, CHIT1, LCN2, OLR1* and *SLC2A).* The hub gene *ELANE* encodes a serine protease secreted by neutrophils that is known to regulate the function of natural killer cells, monocytes and granulocytes and is essential for neutrophils in fighting infections^71,72^. Neutrophil activation is characteristic of severe COVID-19^73^ and shared with other inflammatory states^74^. Module 26, identified as shared between PTB and RSV-LRTI, includes the hub gene *TAP1* which is known for its antiviral activity through Type I interferon production^75^. Other hub genes include *STAT, PSMB8, GBP1, PSMB9, HLA-E, GBP5, HLA-F, GBP2, IRF9, APOL3* and *CASP1* which are also known be associated with COVID-*19*^76^. The detailed enrichment for GO terms are provided in Supplementary Table S5 and Supplementary Fig 4.

Cell proportion estimation showed that in children hospitalised with COVID-19 there was depletion of macrophages and monocytes compared to healthy controls. In contrast, in children hospitalised with RSV-LRTI, increased proportions of regulatory T-cells and macrophages, and a depletion of T-cells and class switched memory B-cells were observed. Similarly, for PTB, there was an increase in macrophages, monocytes and neutrophils, and a depletion of T-cells and CD8+ alpha-beta T-cells, and cytotoxic NK cells (Fig 4). The depletion of T and B cells is a key feature of COVID-19 severity^77^. T-cell immunity is essential to control PTB^78^.

We identified 247 genes that predicted the severity of LRTI. Ten were common among LRTIs. *IL16* is involved in pro-inflammatory responses to activate T-cells and the production of cytokines^79^. Five genes could discriminate hospitalised children with LRTI including: *PITPNC1*, *TPX2*, *LARP1*, *HABP4* and *SMIM10L2A*. *PITPNC1* is known for pulmonary function and asthma^80^. Five genes, including *PAFAH2, LINC02915, CLSPN, EIF4G1* and *IFI27,* were predictors of both COVID-19 and RSV-LRTI hospitalization. *PAFAH2* is known to be associated with pulmonary micro-thromboses linked with LRTI severity^81,82^. The top COVID-19 predictor, *RORC,* is a key regulator of cellular differentiation, immunity and glucose metabolism. *CBX7* is part of the Polycomb complex required for transcriptional repression of many genes and cancer progression^83^ and is functionally linked with lymphocyte, monocyte and neutrophil counts. *ZFVE9* is known to be predictive of active TB^84^. The *ADAMTS2* is metalloprotease that processes extracellular matrix is implicated in tissue damage^85^ and is a marker for COVID-19 severity across disease conditions^86^.

Assessing the potential druggability of differentially expressed genes can help in prioritizing drug targets. Amongst the DEGs for LRTIs, known approved drug targets (TClin) were identified including: *KCND3, CACNA1E, GABRG2, CHRNA5, KCND1* and *ADRB2* for severe COVID-19; *GABRG2, KCND1, CA12, CACNA1A, IMPDH2* and *PDE1B* for RSV-LRTI; *CACNA1E, GABRG2, KCNK3, CHRNA5* and *CHRNB2* for PTB as shown in Supplementary Table S8. Interestingly, the top predictors of severity were not previously identified as drug targets, including CBX*7, MYBL1, VRK3, ZNF324, KRTAP5-1* and *GPR153.* In the top PTB predictors, *NR3C2* and *GPR15* have high scores for Tclin but the top predictors, *MID2* and *ZFYVE9,* have not previously been identified as drug targets showing opportunity for drug target prioritization for this population. For RSV-LRTI, except for *RORC,* most top predictors (*IFI27, CBX7, PNRC1* and *IGF2BP3)* have not previously been targeted for drug development (Supplementary Table S9).

One of the strengths of our study is the assessment of hospitalised children with one of the three major LRTIs in children in LMICs and comparison with healthy children using datasets generated from a similar genetic background. Many known signature-genes identified for COVID-19 (*IFI27, OLFM4*), RSV-LRTI (*SIGLEC1, ISG15, IFI44)* and PTB (*MMP8, MMP9, DEFA1, DEFA1B, DEFA3* and *DEFA4*) are known to be associated with progressive severity^87^, showing the reproducibility of our findings. A limitation is that the DCHS children were older than children with LRTI, but we used age as a covariate to overcome this confounding effect.

From our transcriptomic analysis of children with LRTIs due to three different aetiologies, we have identified novel data providing key immune response related genes associated with severity for children hospitalised with COVID-19, RSV-LRTI and PTB in African children. These genes can be used for baseline characterization, as predictive markers for respiratory infection severity and as potential therapeutic targets.

## Data availability

Supplementary data and summary statistics for transcriptome wide association analyses are available from: DOI https://doi.org/10.5258/SOTON/D3587.

An anonymised, de-identified version with data can be made available on request. All requests should be directed to Prof Heather Zar, DCHS Study Principal Investigator.

## Code availability

The custom code used to generate graphics is available at GitHub repository: https://github.com/negusse2025/respiratory_infections.git.

## Acknowledgements

Funding, participants/ families and staff. The some of the graphic illustrations for graphic abstract were accessed from NIH BIOART, including TB (https://bioart.niaid.nih.gov/bioart/527), child (https://bioart.niaid.nih.gov/bioart/75) and SARS-CoV-2 (https://bioart.niaid.nih.gov/bioart/464).

## Author information

H.J.Z., M.P.K., M.B., N.T.K., D.B. and J.W.H. contributed to conceptualisation.

N.T.K., L.W, H.J.Z. and J.W.H. performed data curation.

N.T.K., E.K. and J.W.H. carried out formal analysis.

N.T.K., M.J, E.K, M.B, D.G, M.P.K. and J.W.H. provided methodology.

H.J.Z performed project administration. N.T.K., H.J.Z. and J.W.H. performed writing—original draft. N.T.K., C.C., D.B., E.K., L.W., M.B., M.J., M.P.K., H.J.Z. and J.W.H. contributed to writing—review, editing and final approval.

## Supplementary data

Supplementary Fig 1 Module dendrogram

Supplementary Fig 2 Distribution of genes per module

Supplementary Fig 3 Venn diagram shared modules between respiratory infections

Supplementary Fig 4 REVIGO biological process for correlated modules with LRTI

Supplementary Fig 5 Blood composition comparisons between LRTI

Supplementary Fig 6 Shared cell types

Supplementary Fig 7 Shared severity predictors for LRTI

Supplementary table S1 Respiratory infection TWAS FDR< 0.05

Supplementary table S2 Respiratory infection TWAS enrichment

Supplementary table S3 WGCNA module genes and eigengene

Supplementary table S4 Network degree distribution for module correlated to LRTI

Supplementary table S5 Enrichment for module correlated to LRTI

Supplementary Table S6. Cell type proportions

Supplementary table S7 severity predictors

Supplementary table S8 Drug target look-up

Supplementary table S9 Severity predictors and target prioritization

Supplementary file Modules_Network_analysis.cys.

---

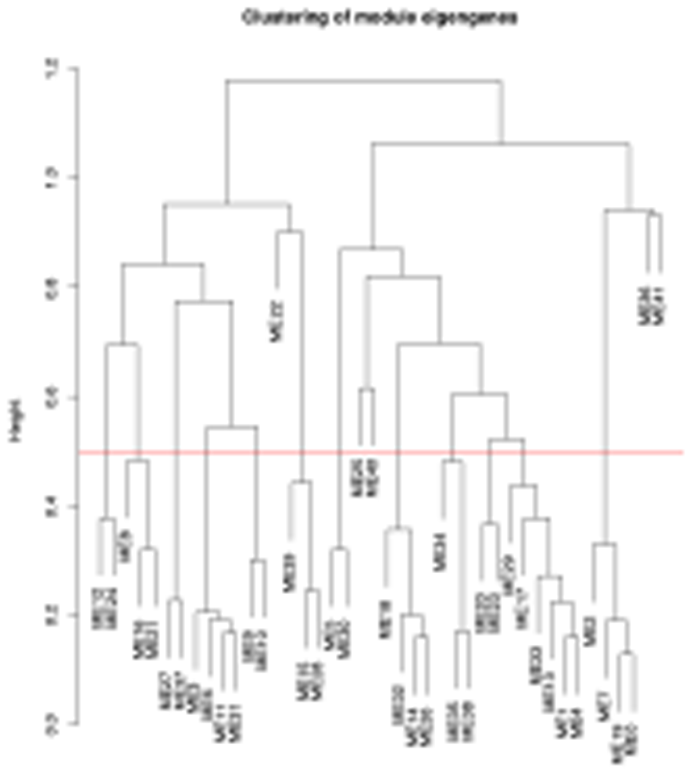

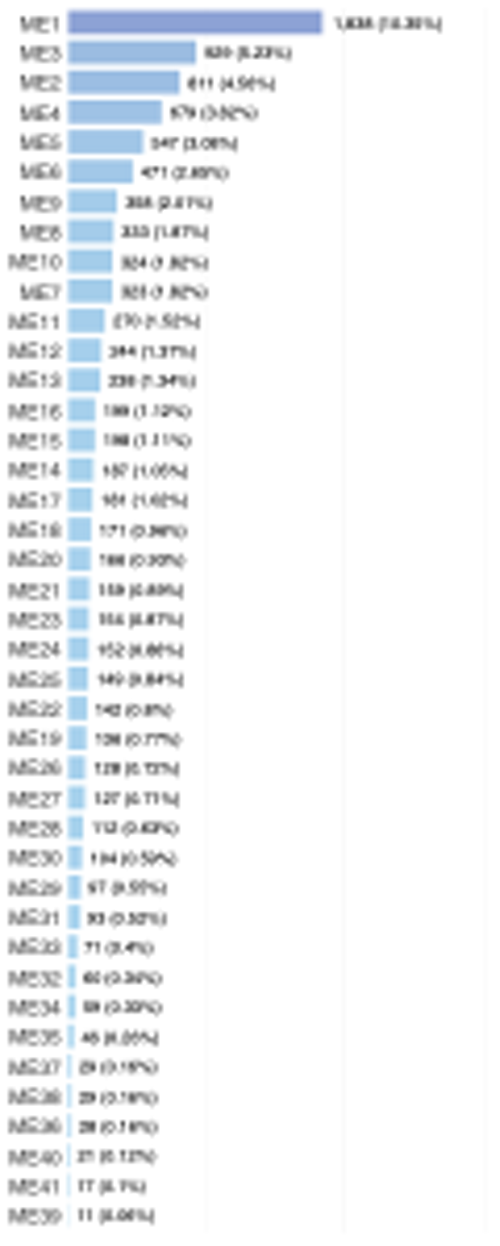

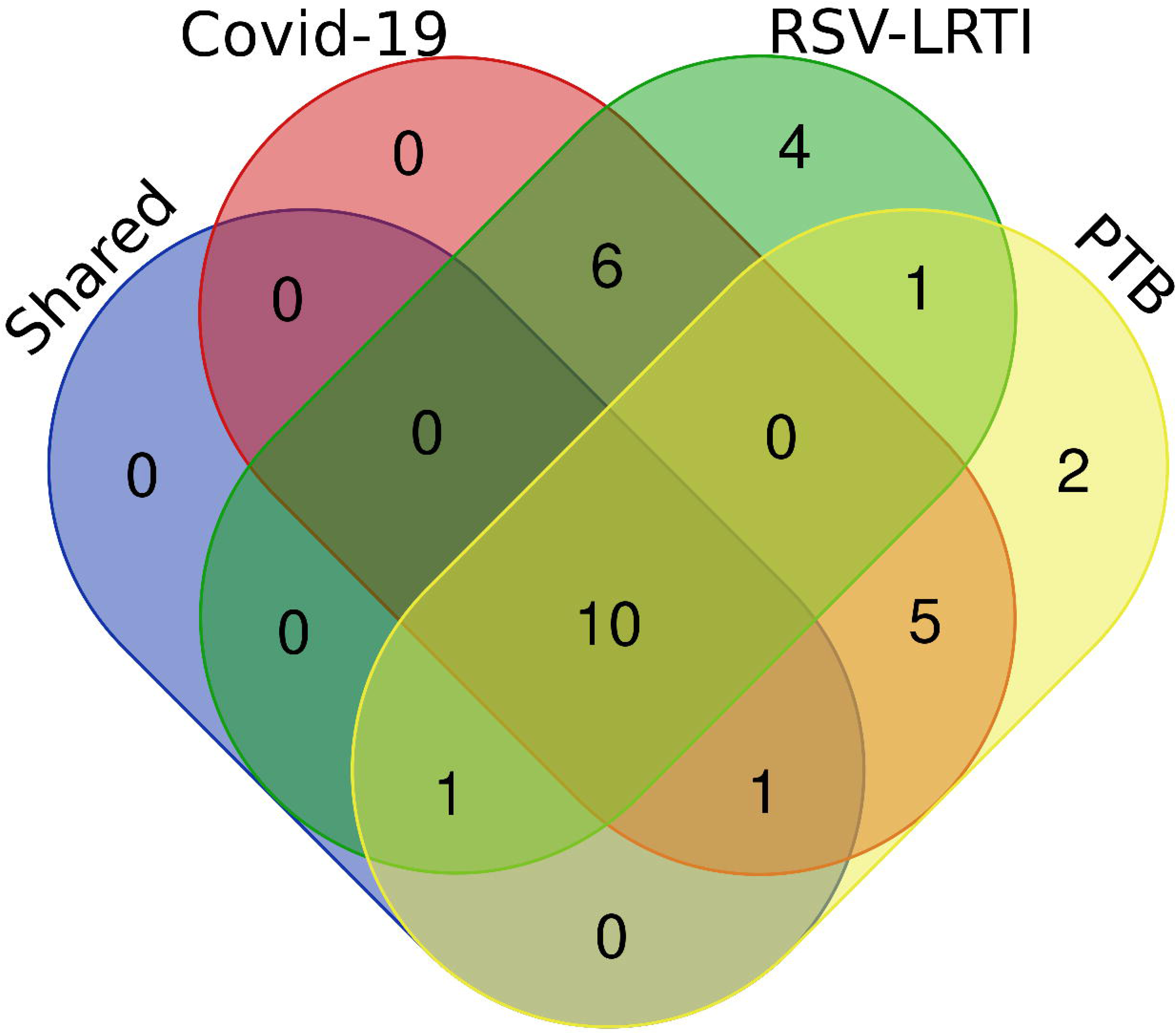

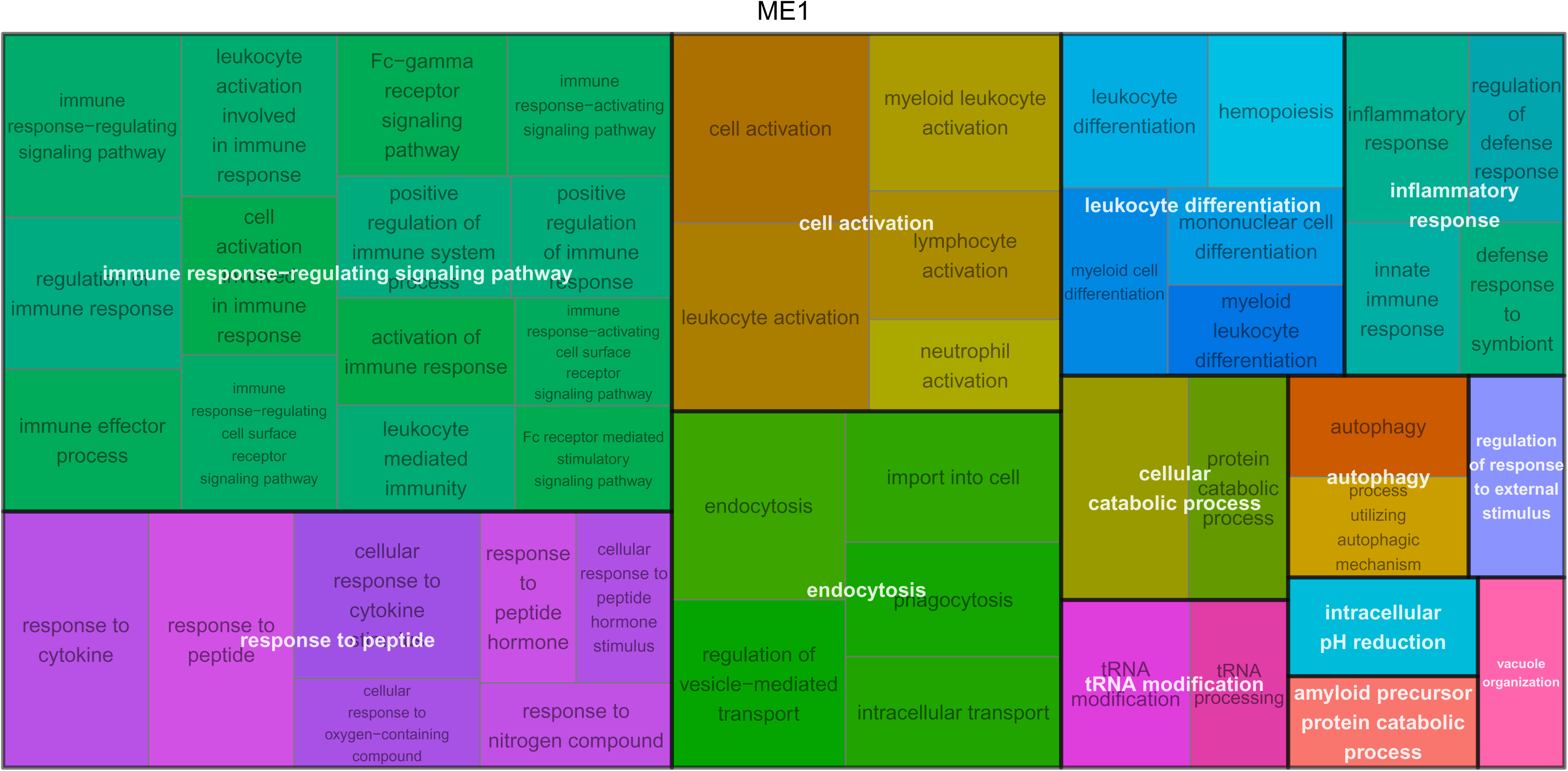

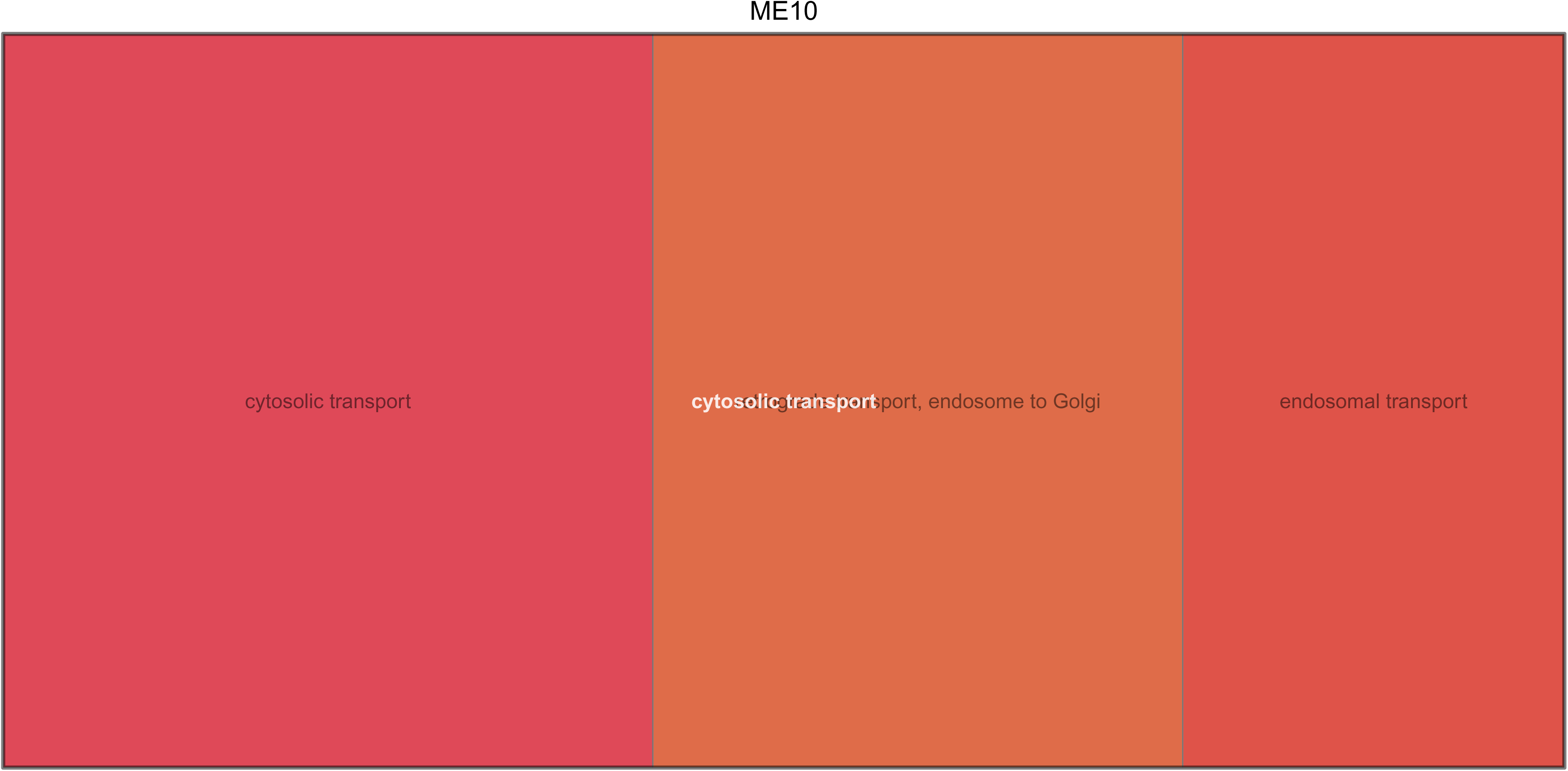

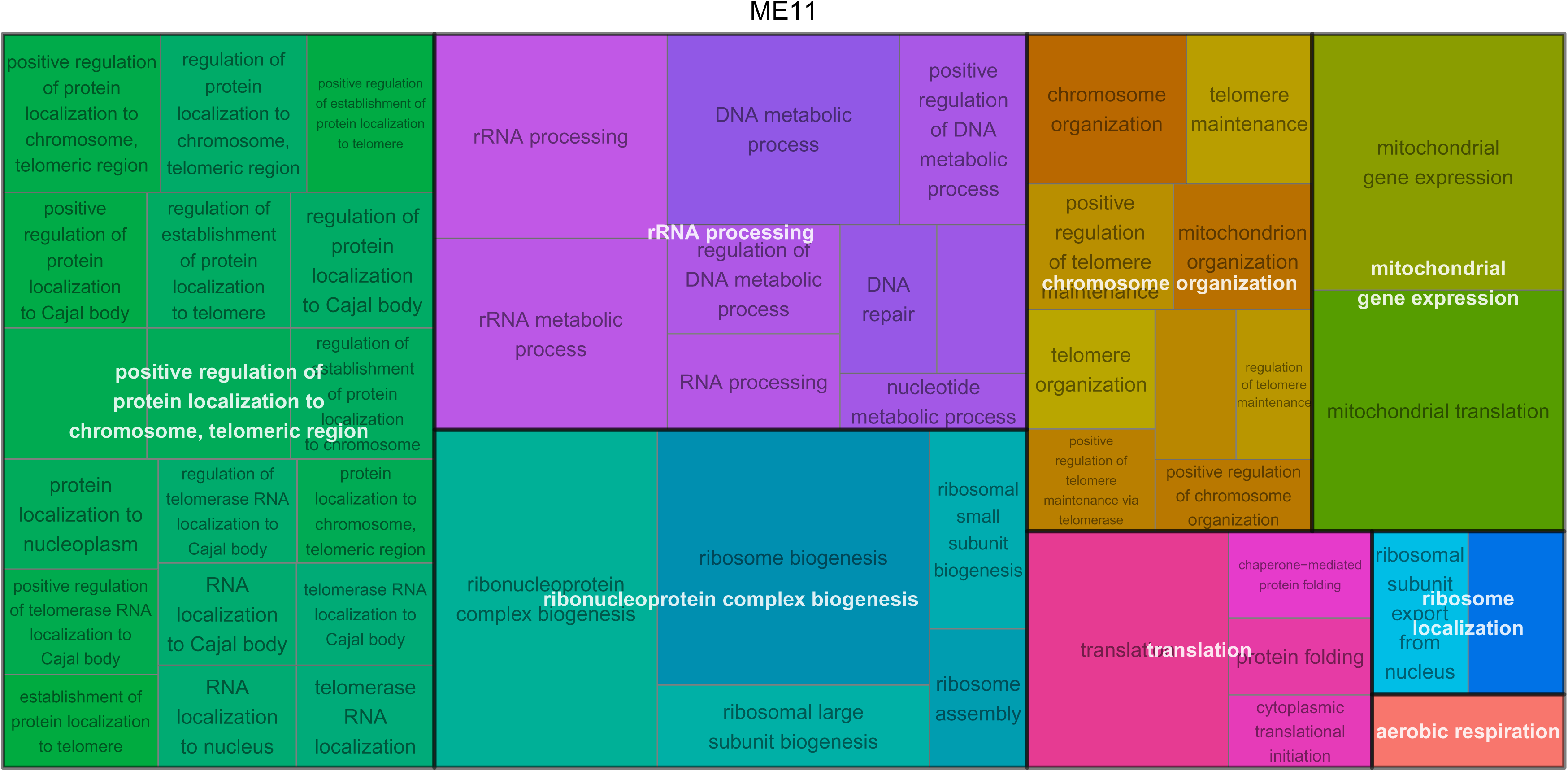

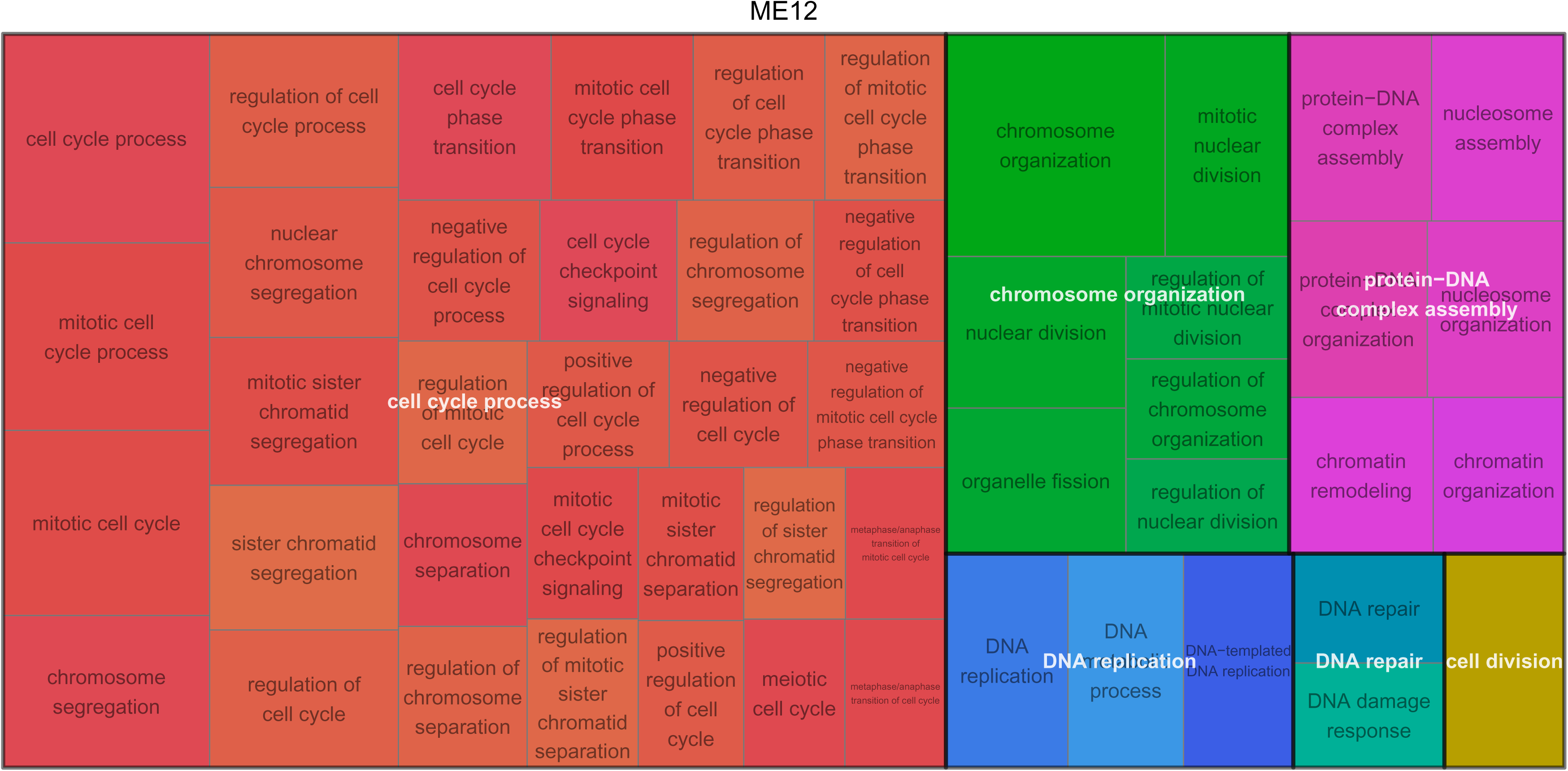

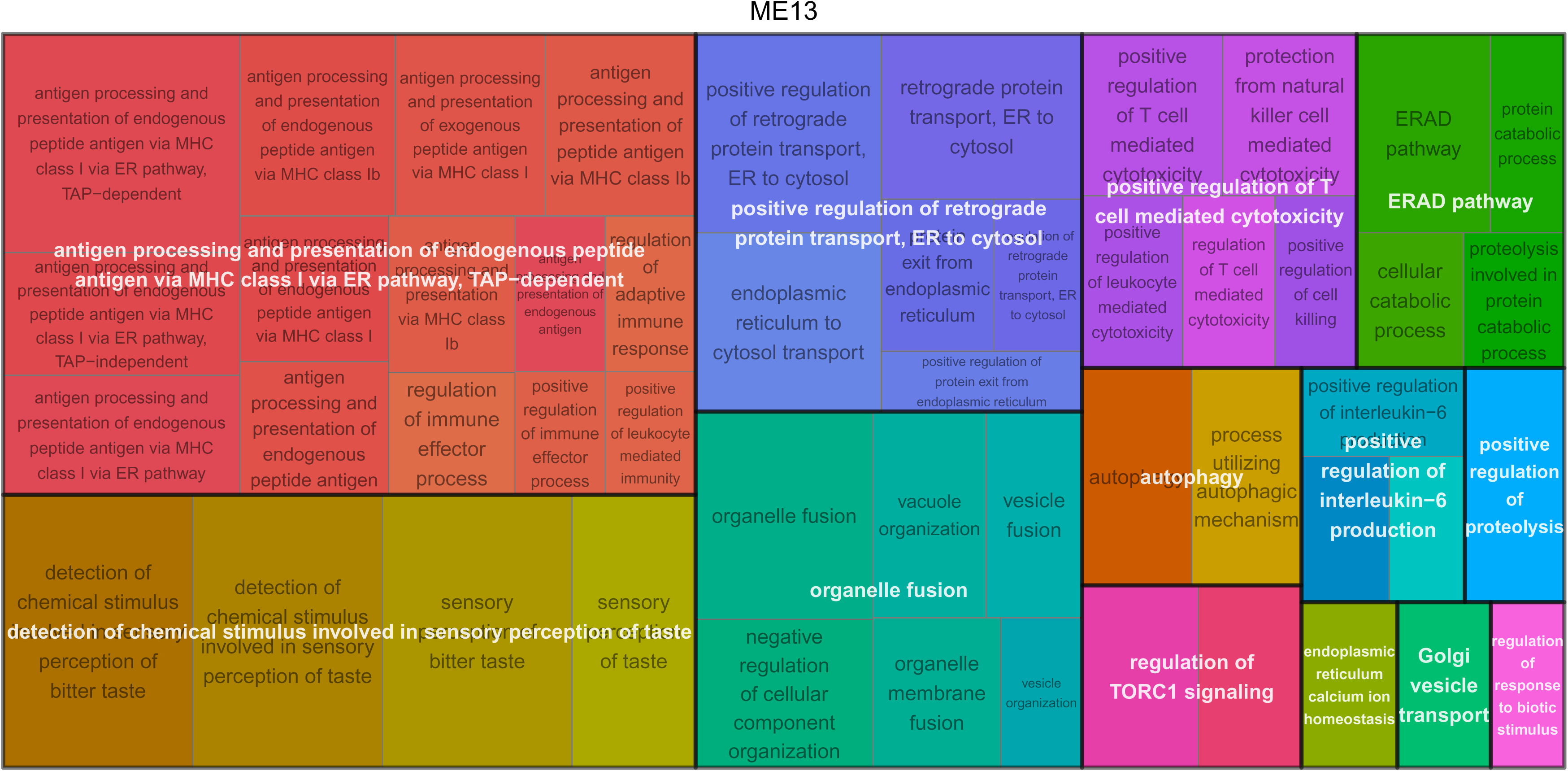

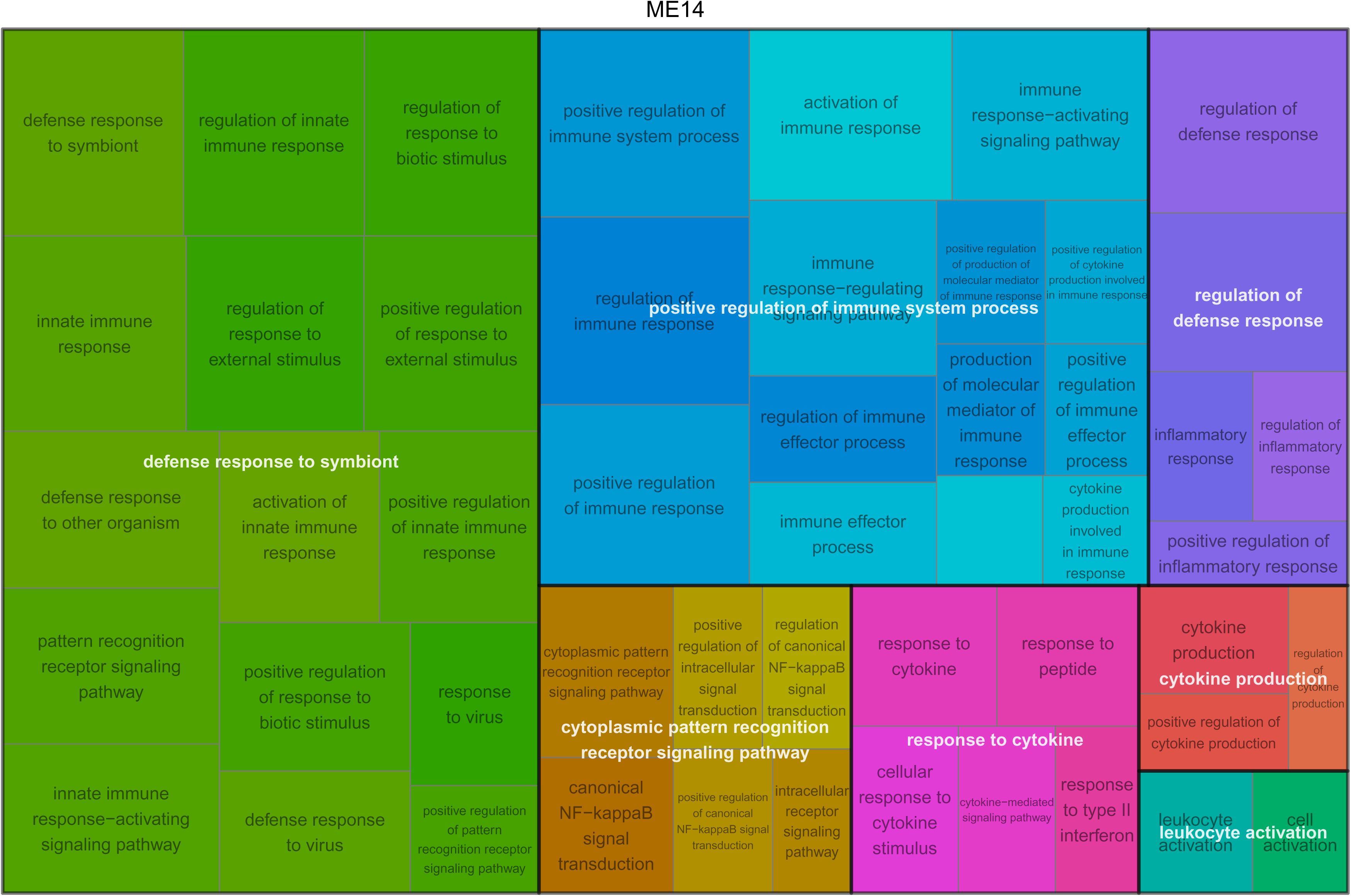

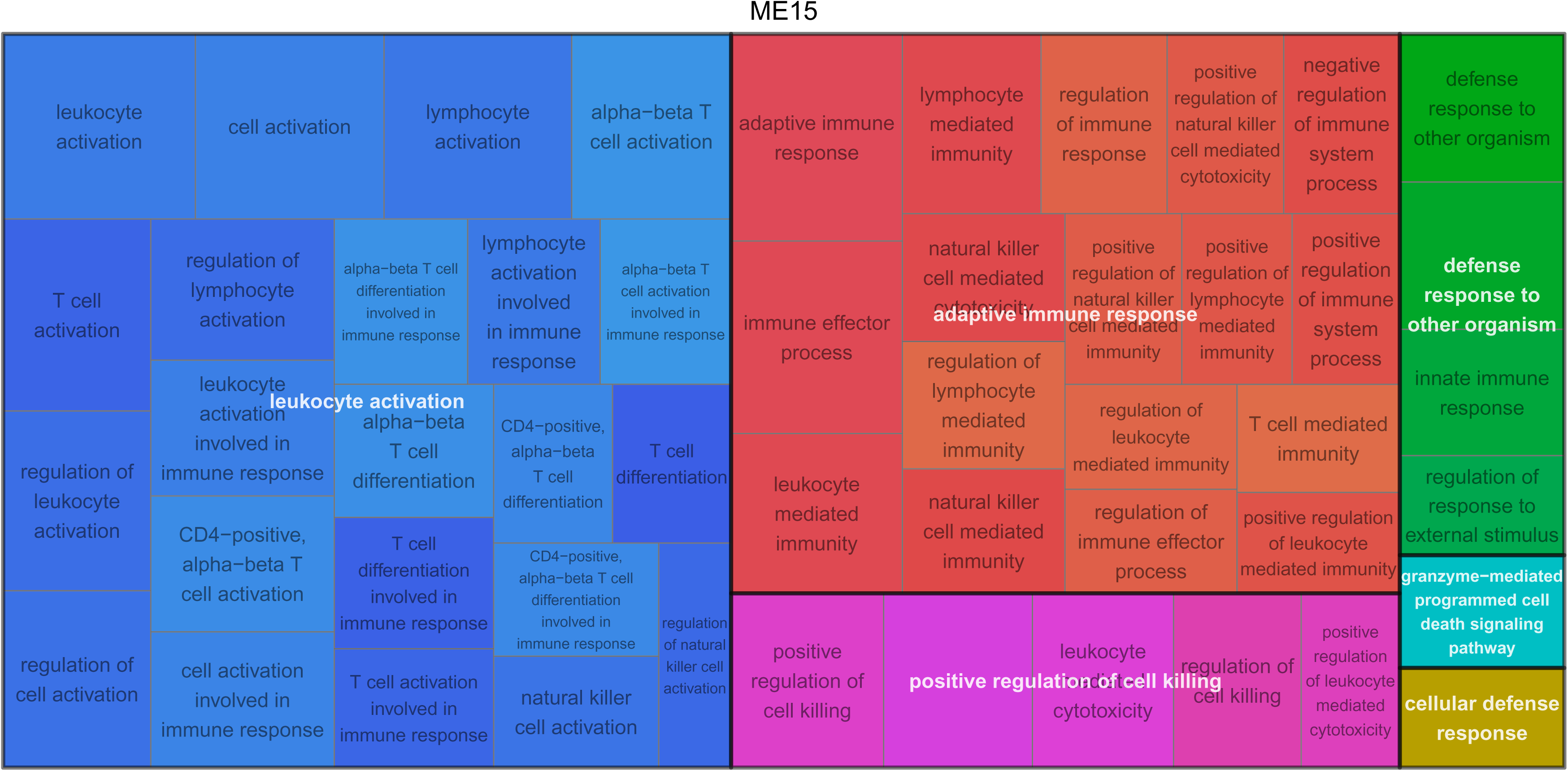

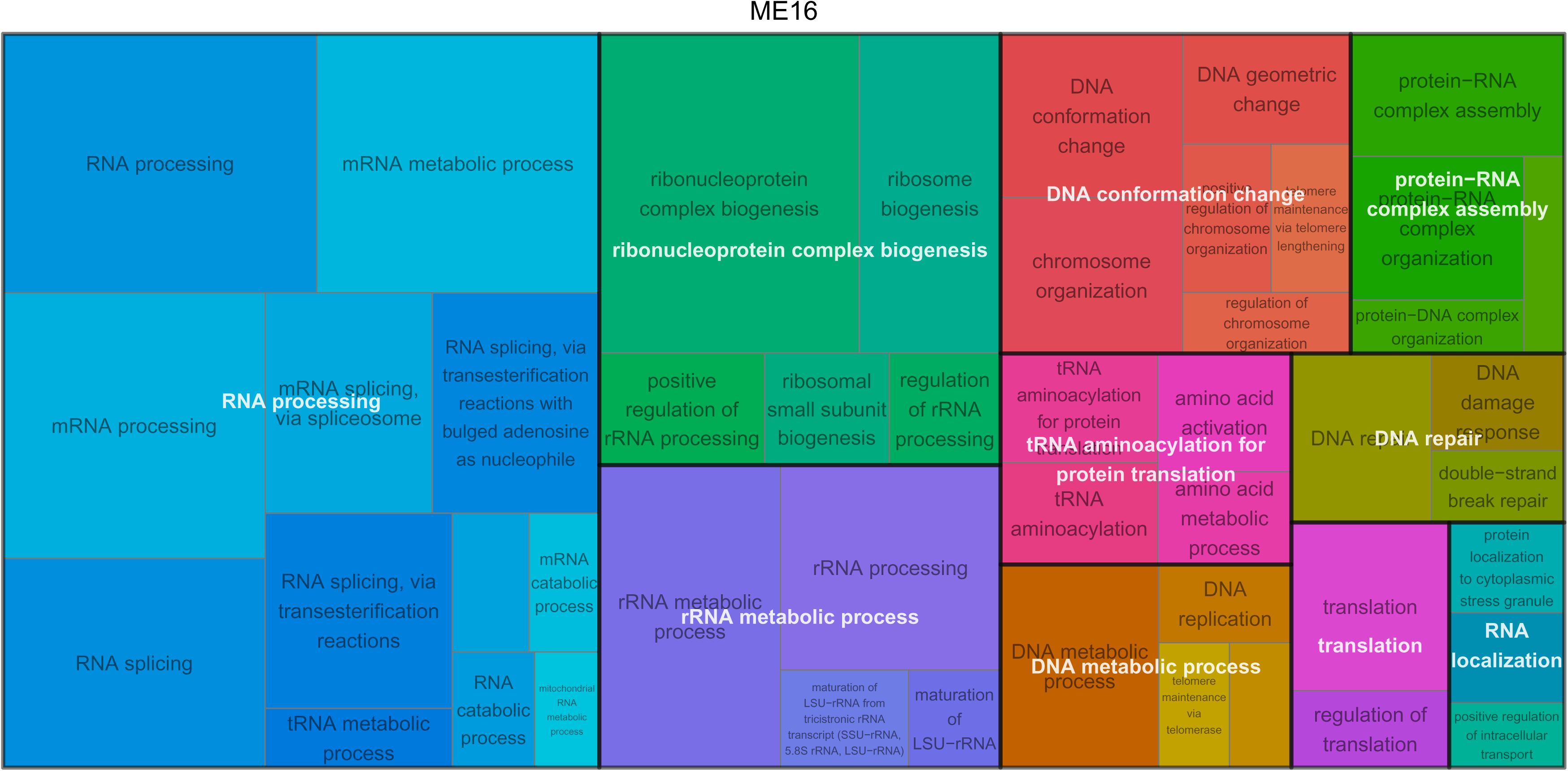

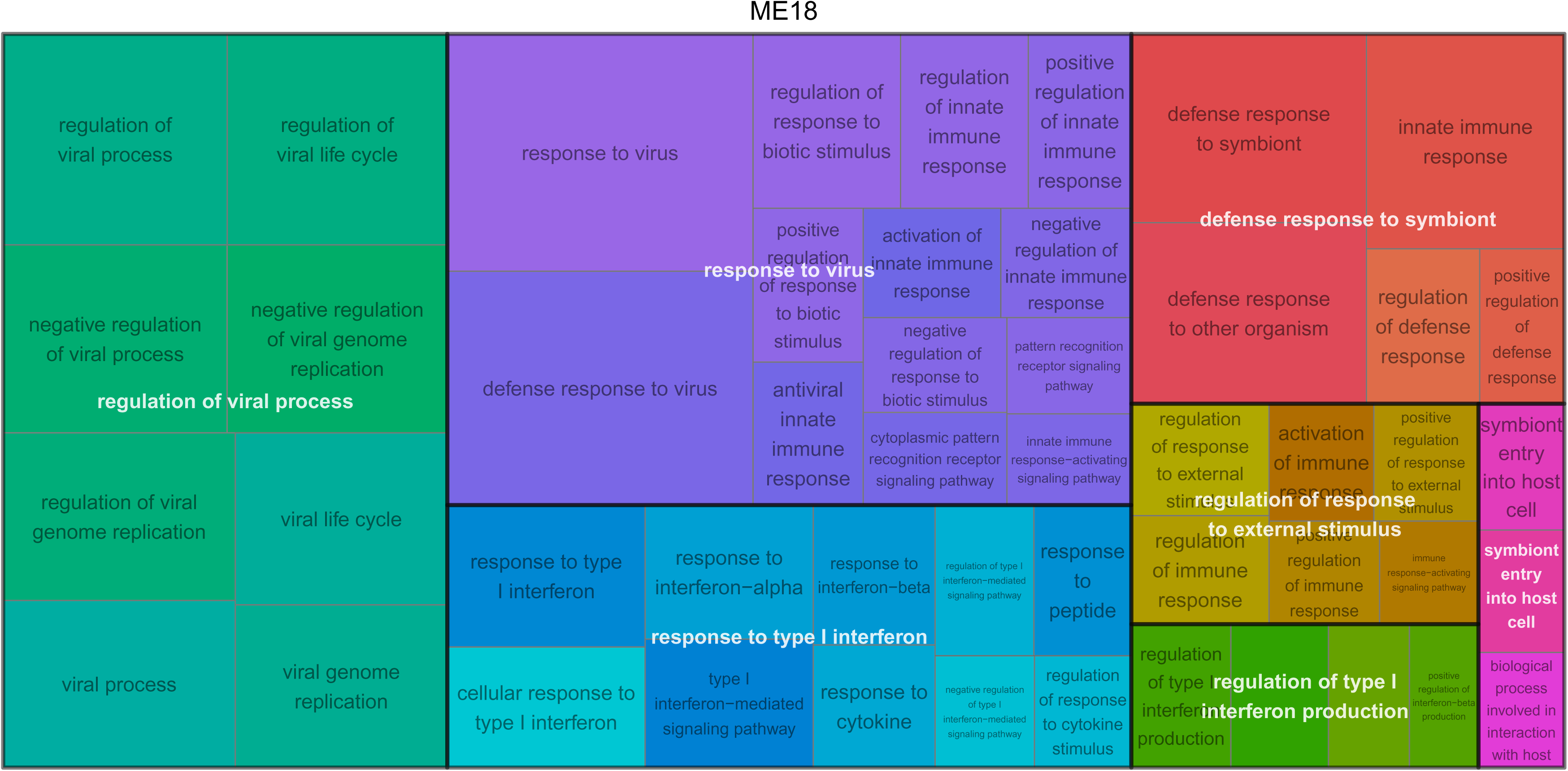

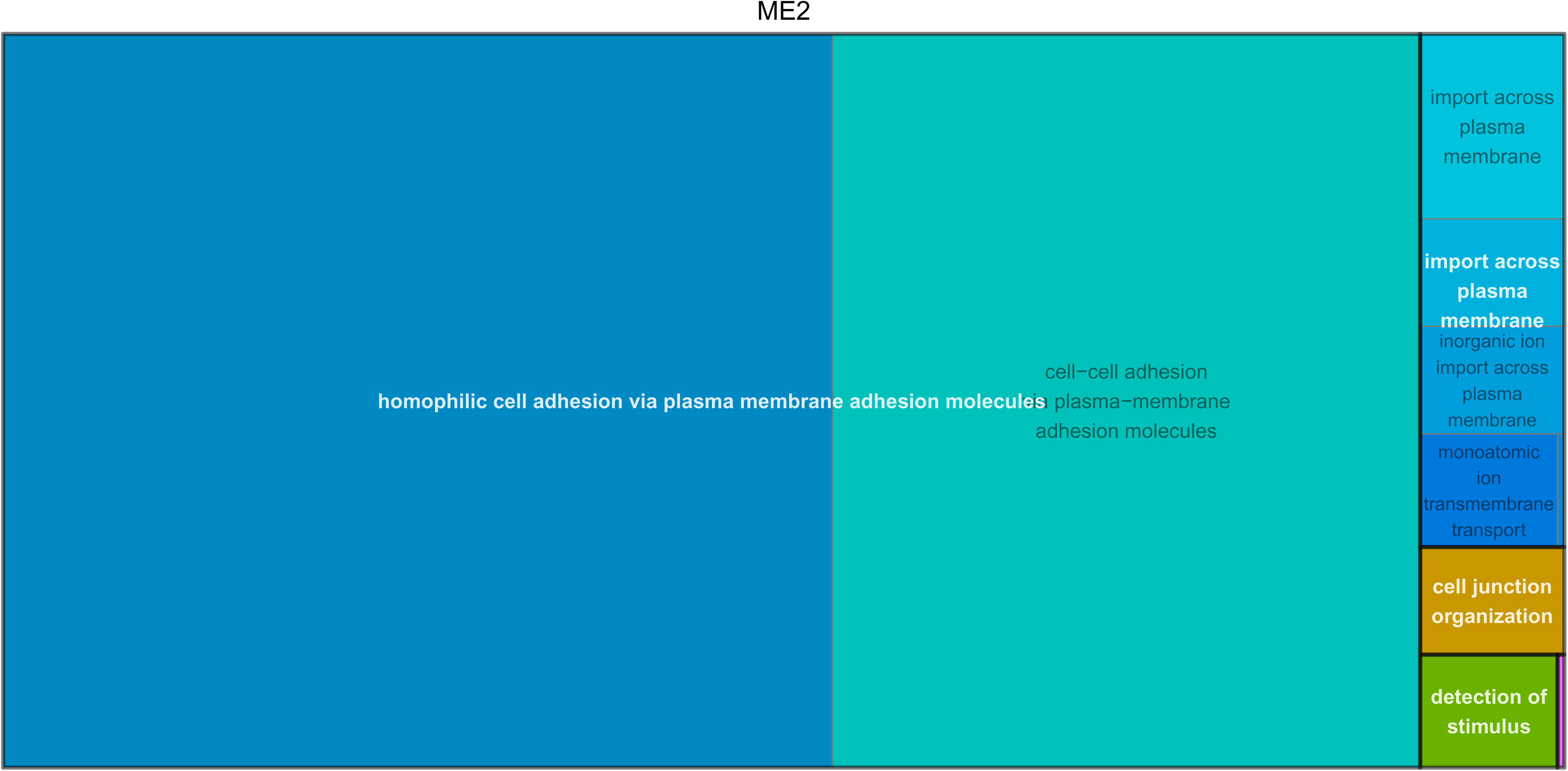

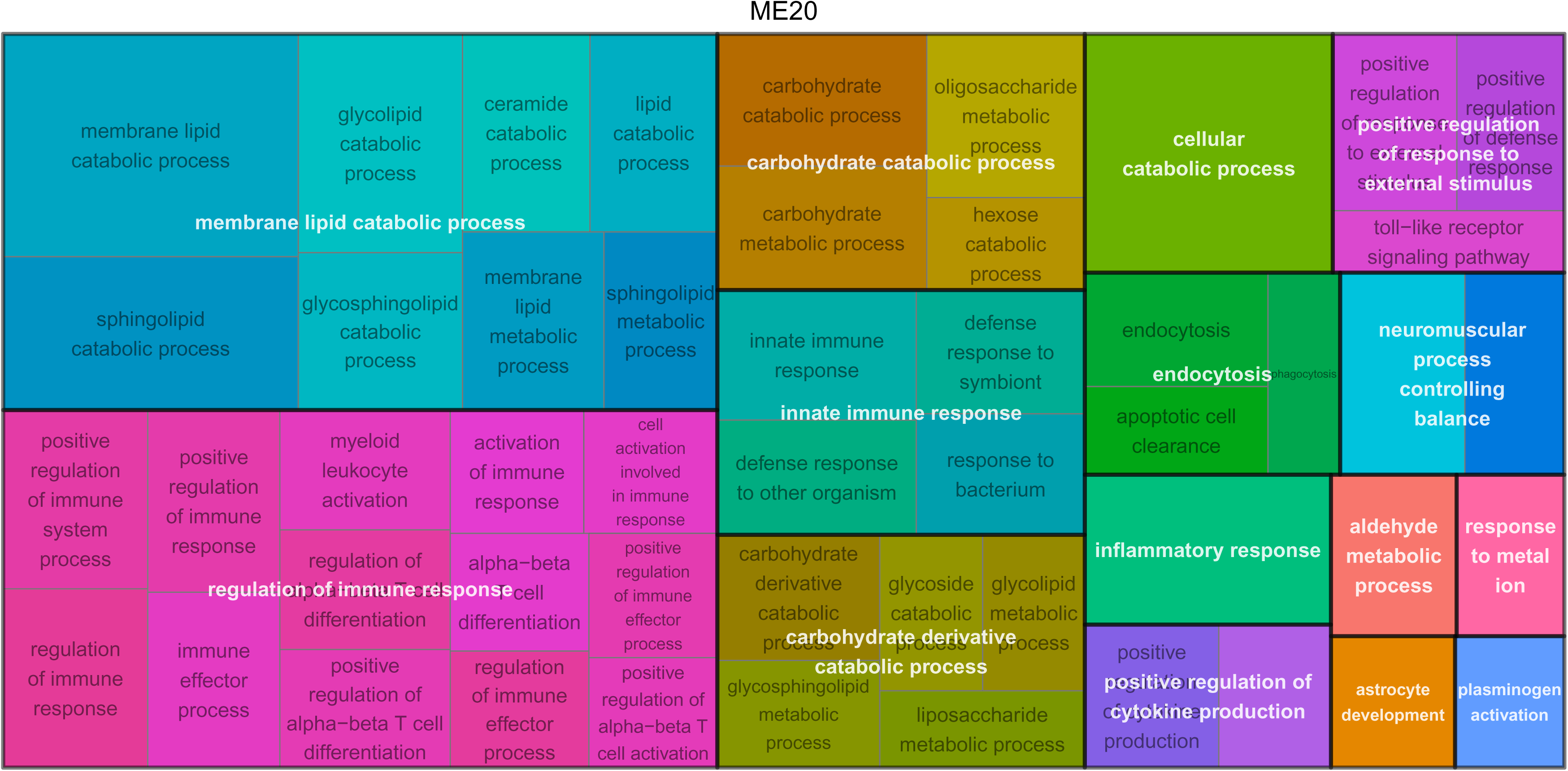

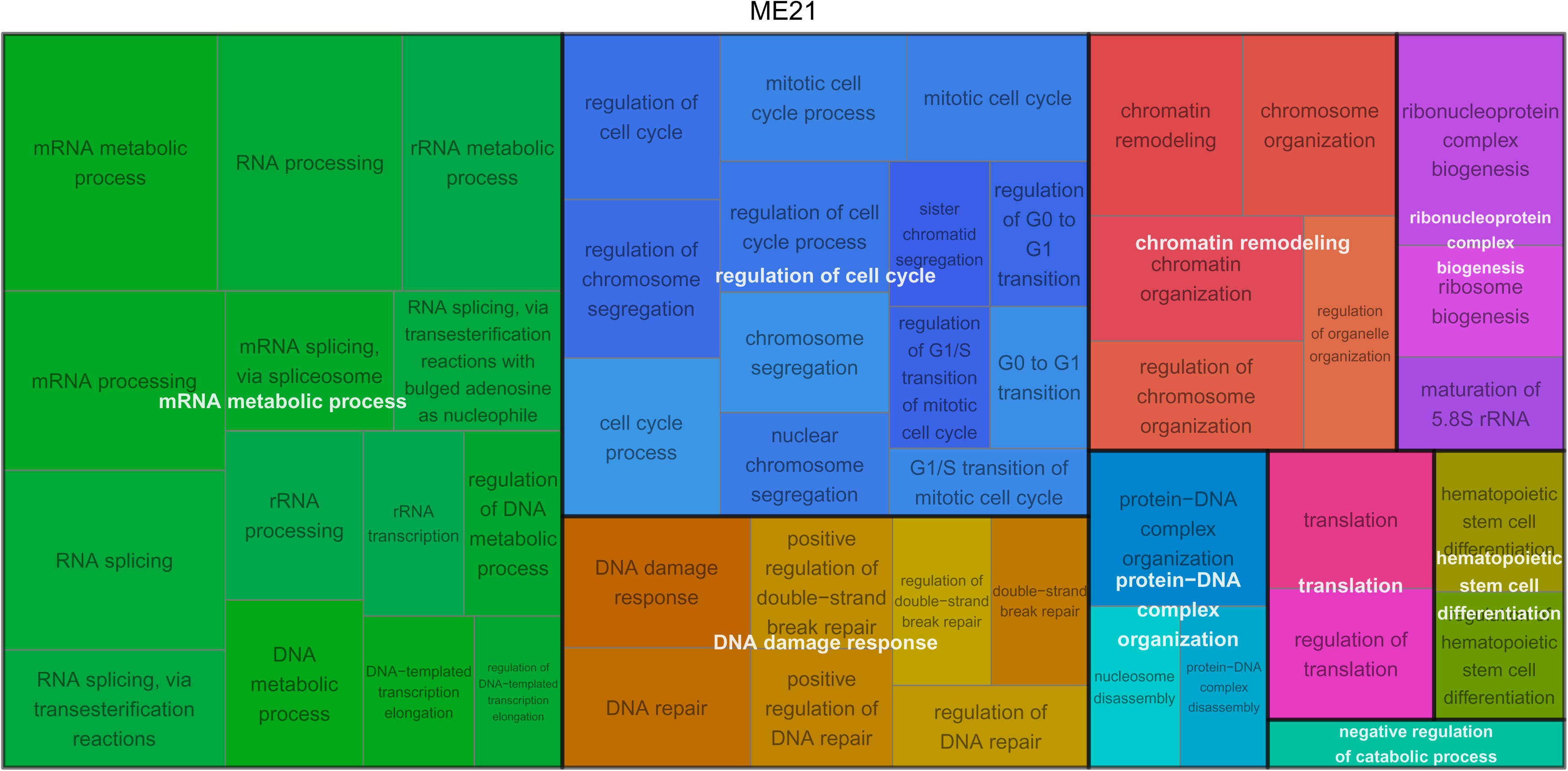

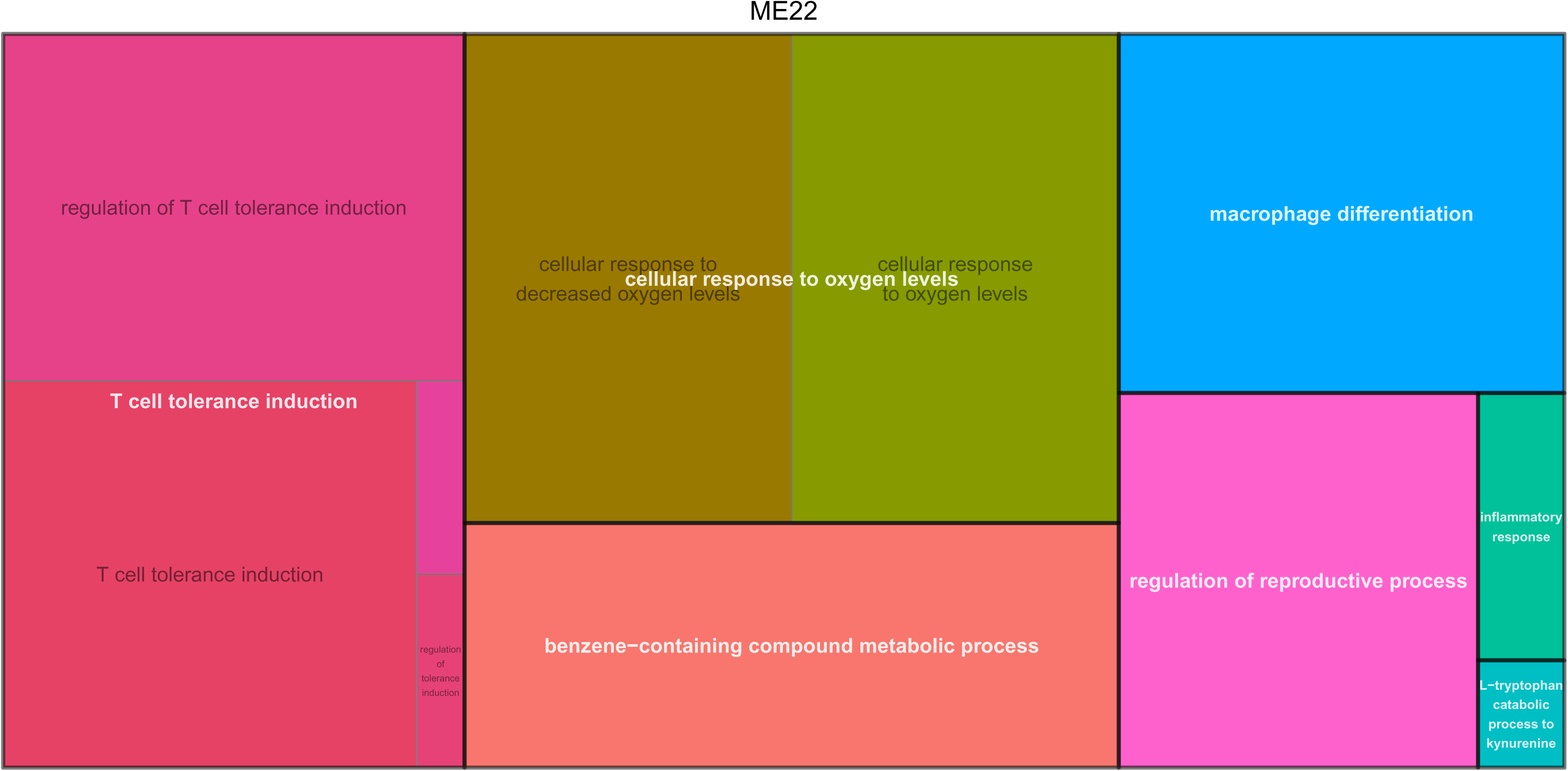

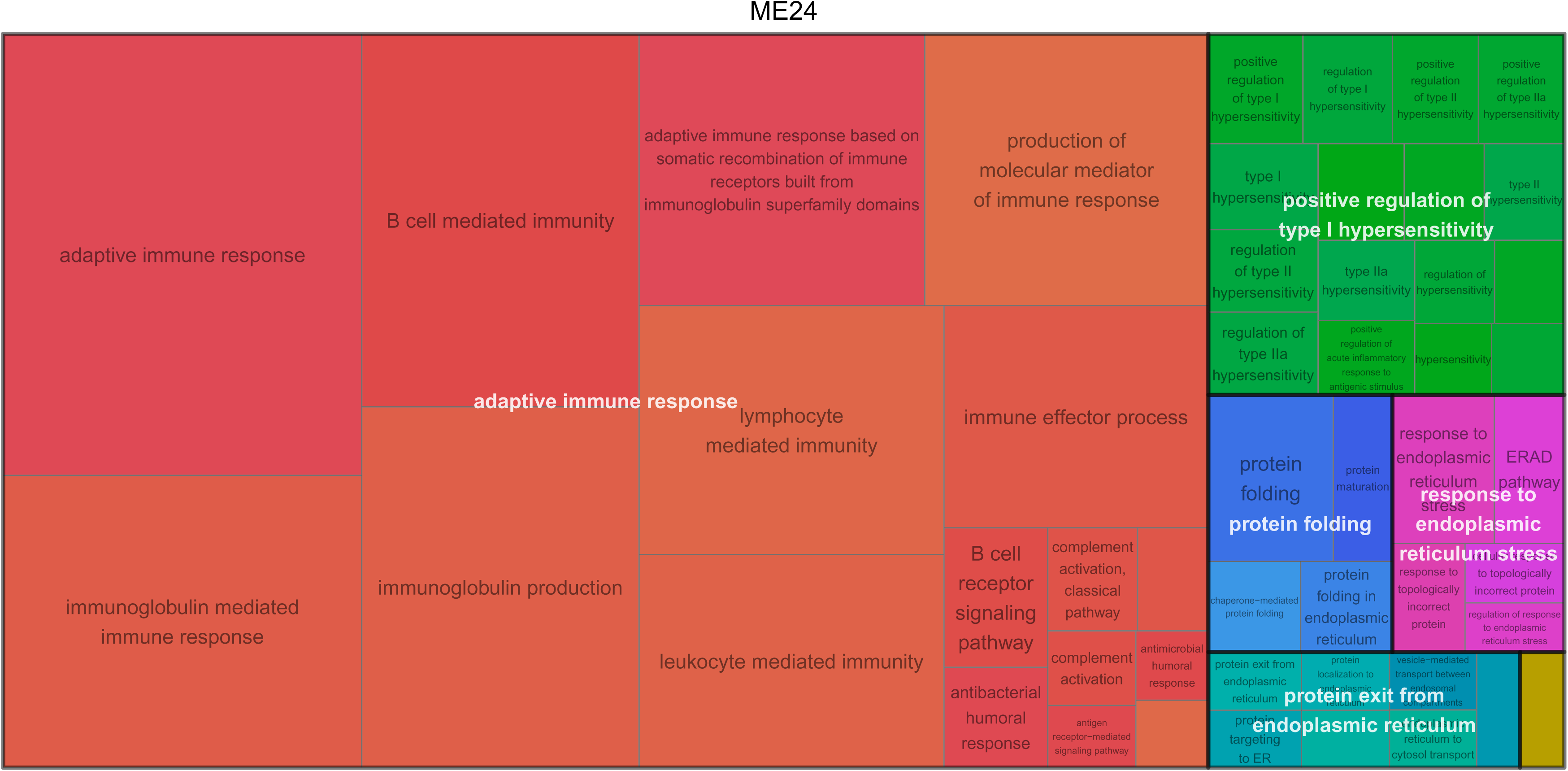

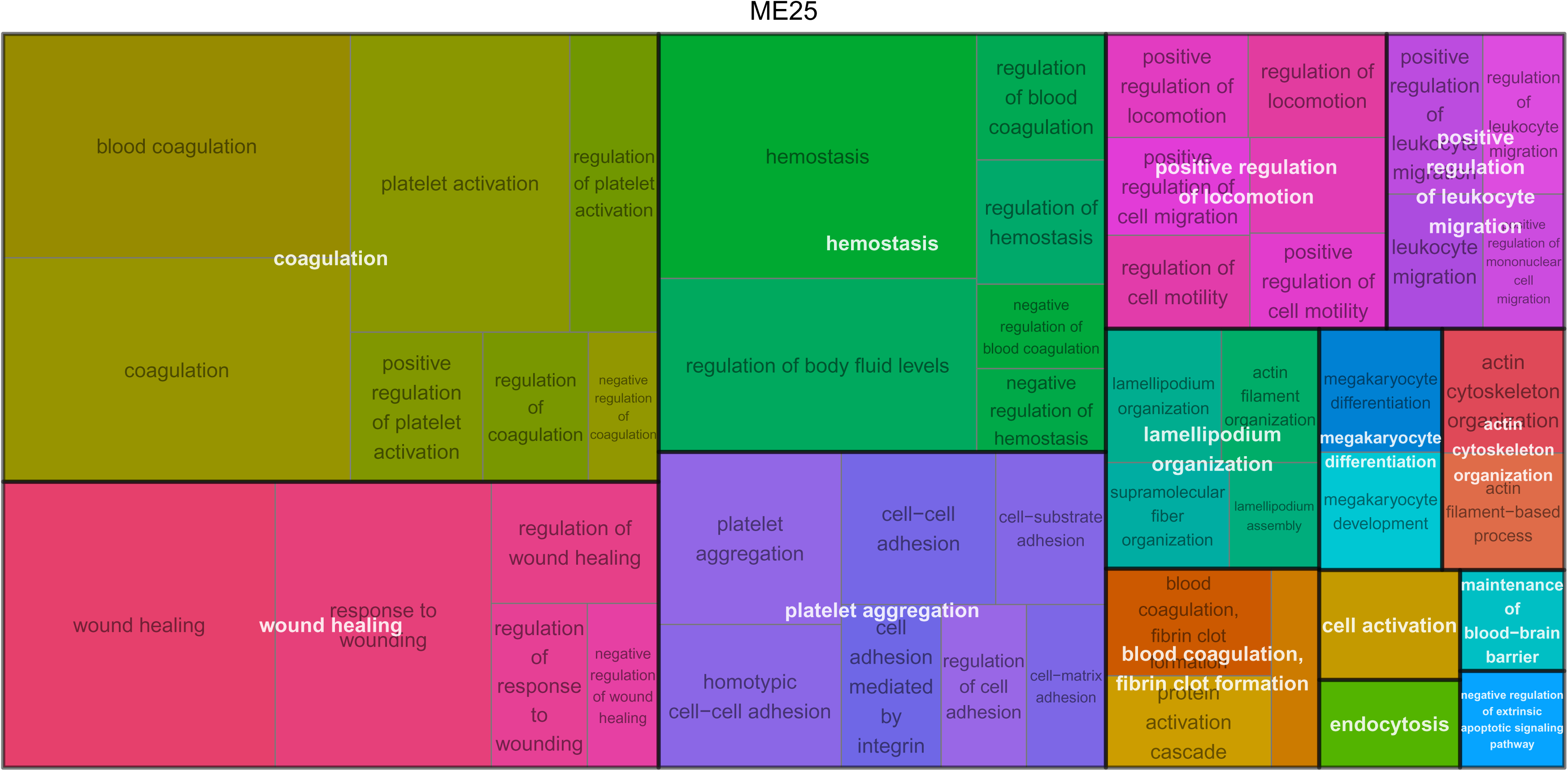

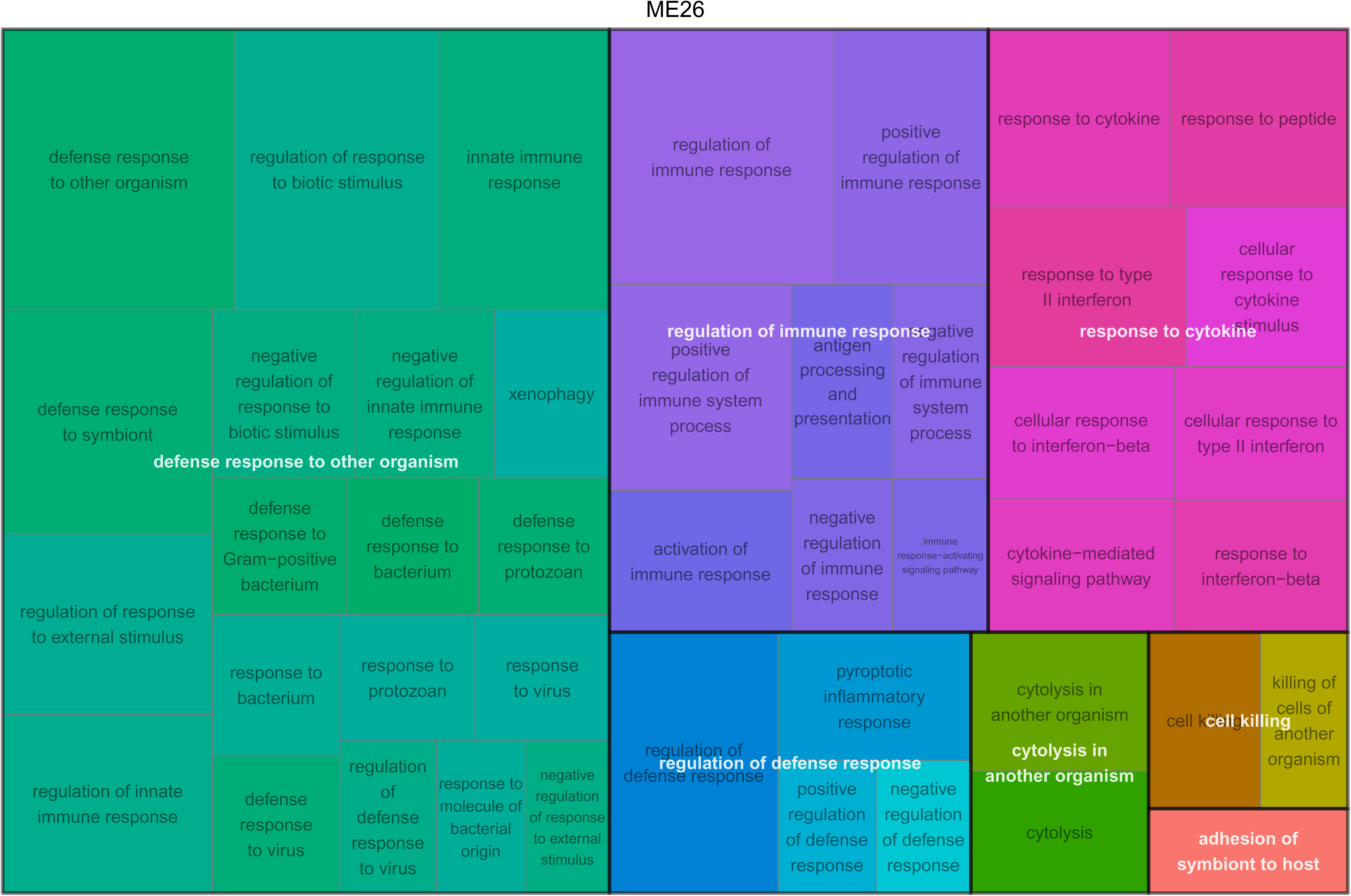

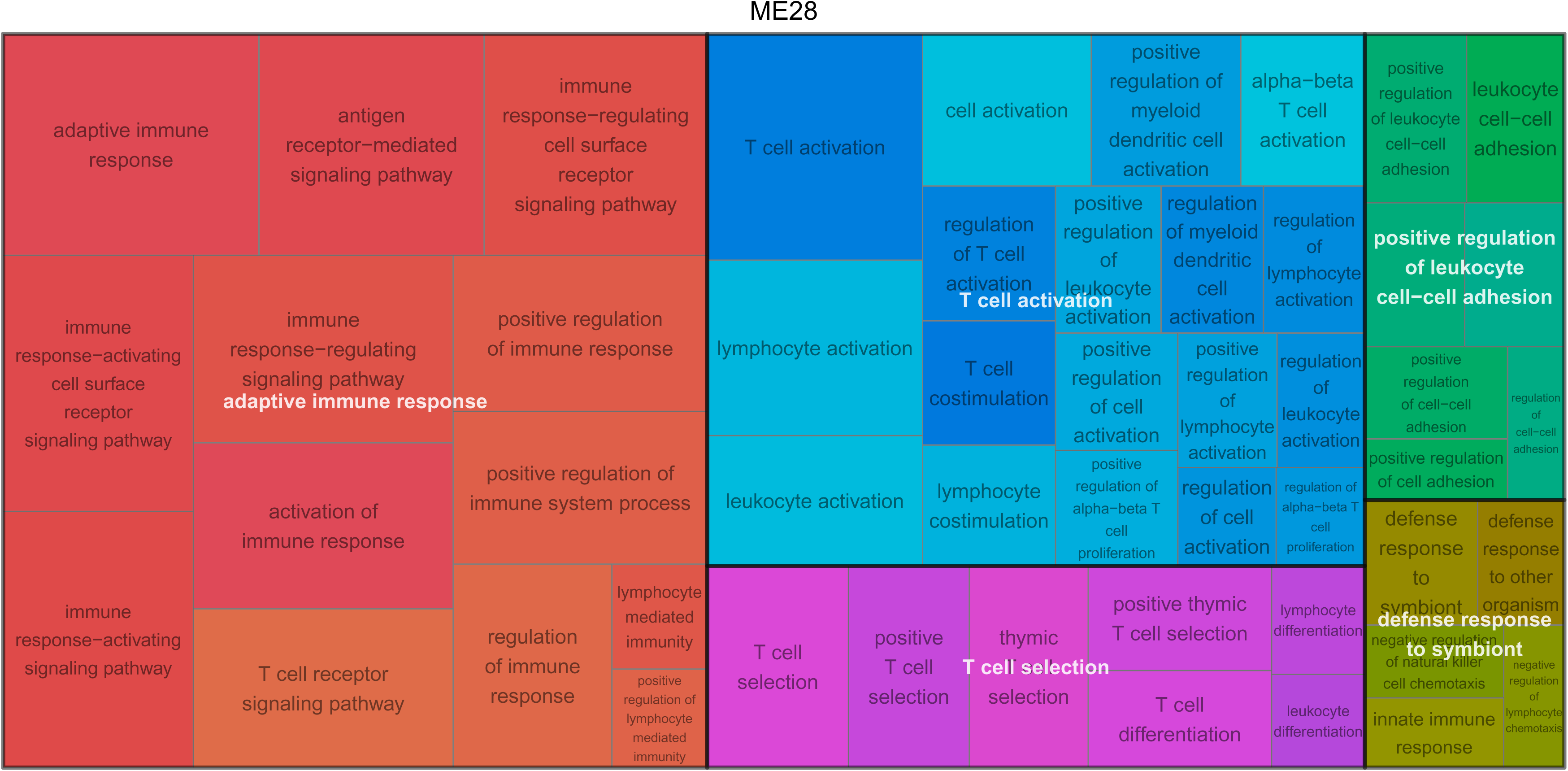

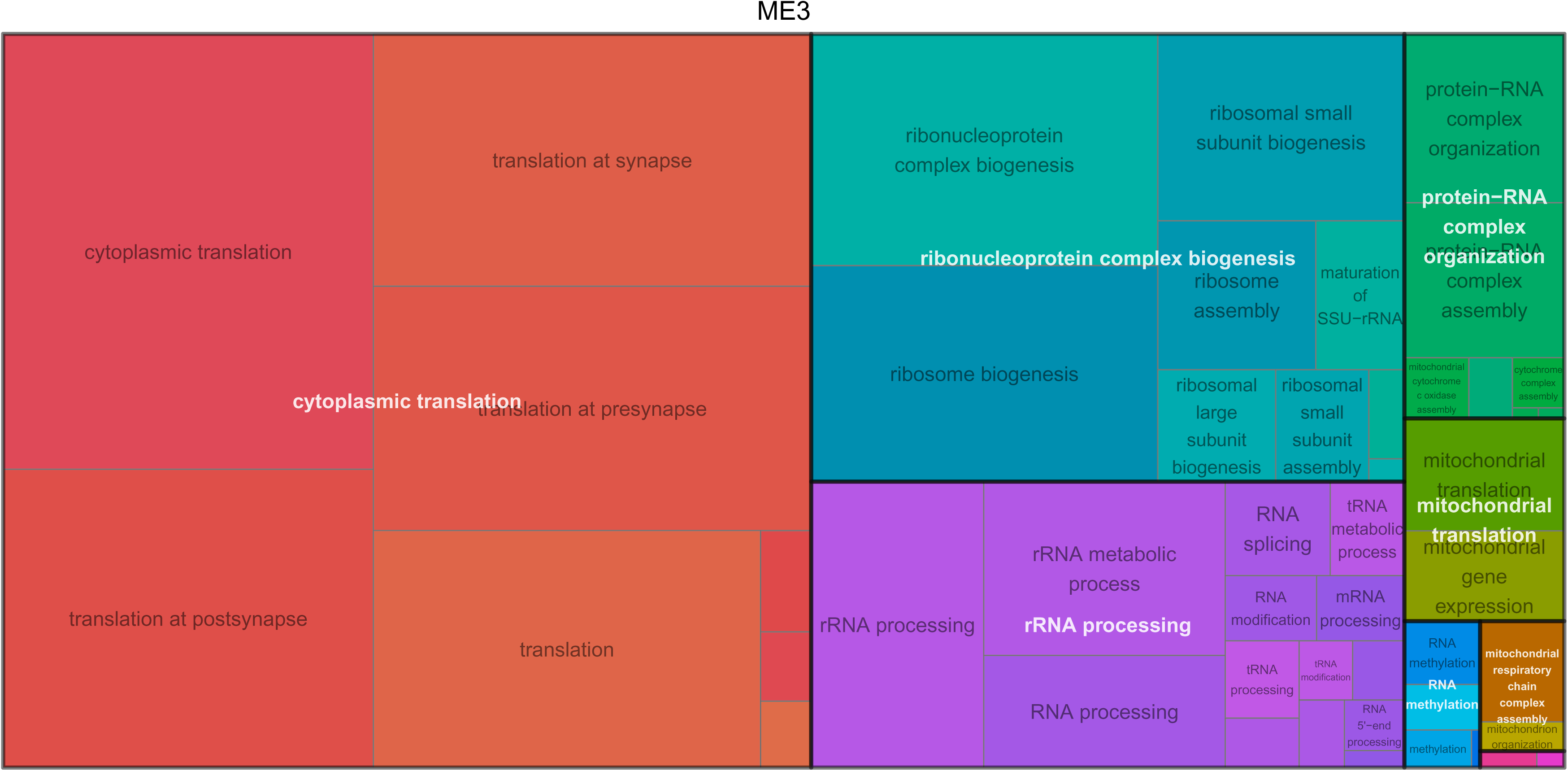

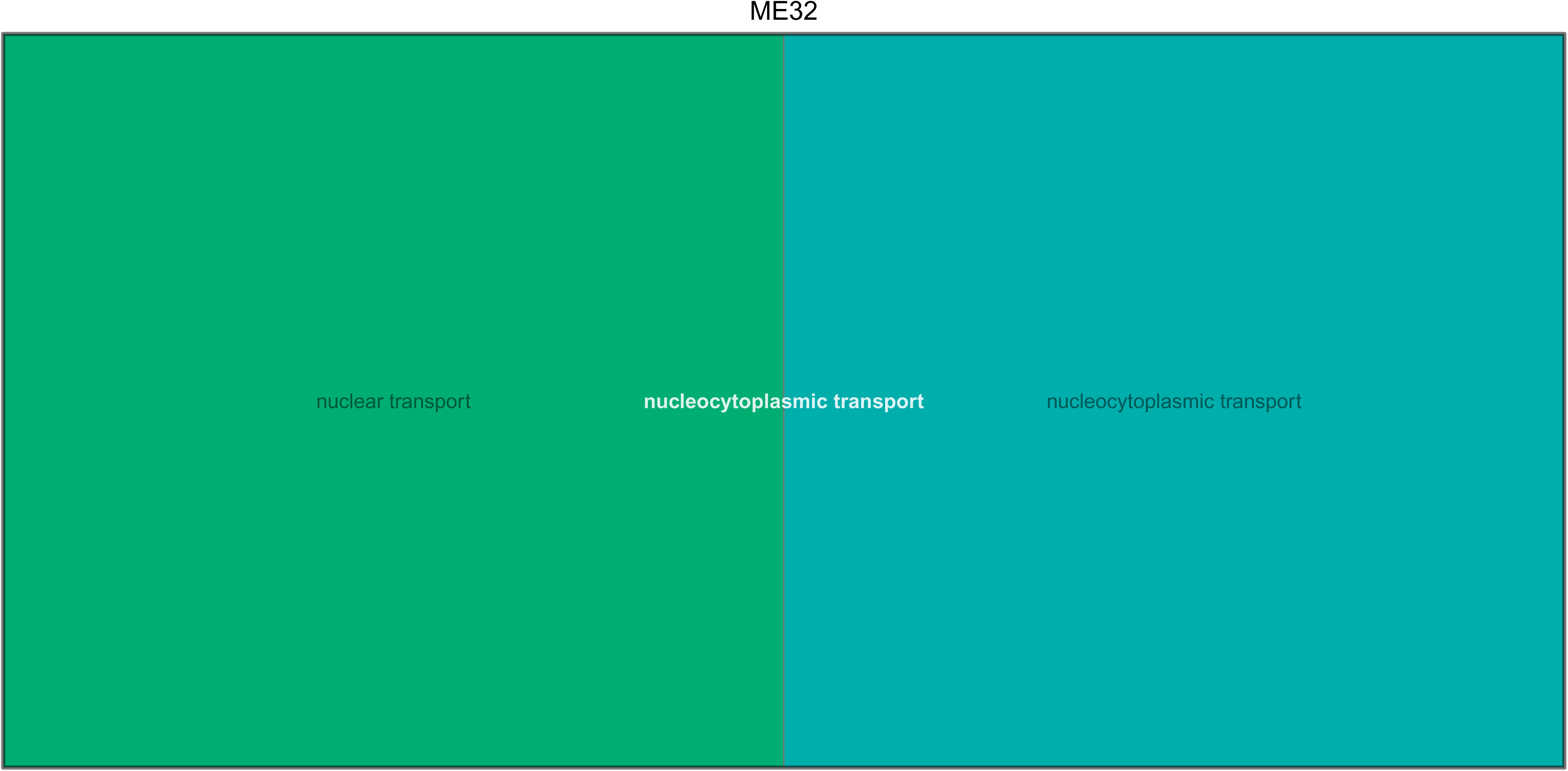

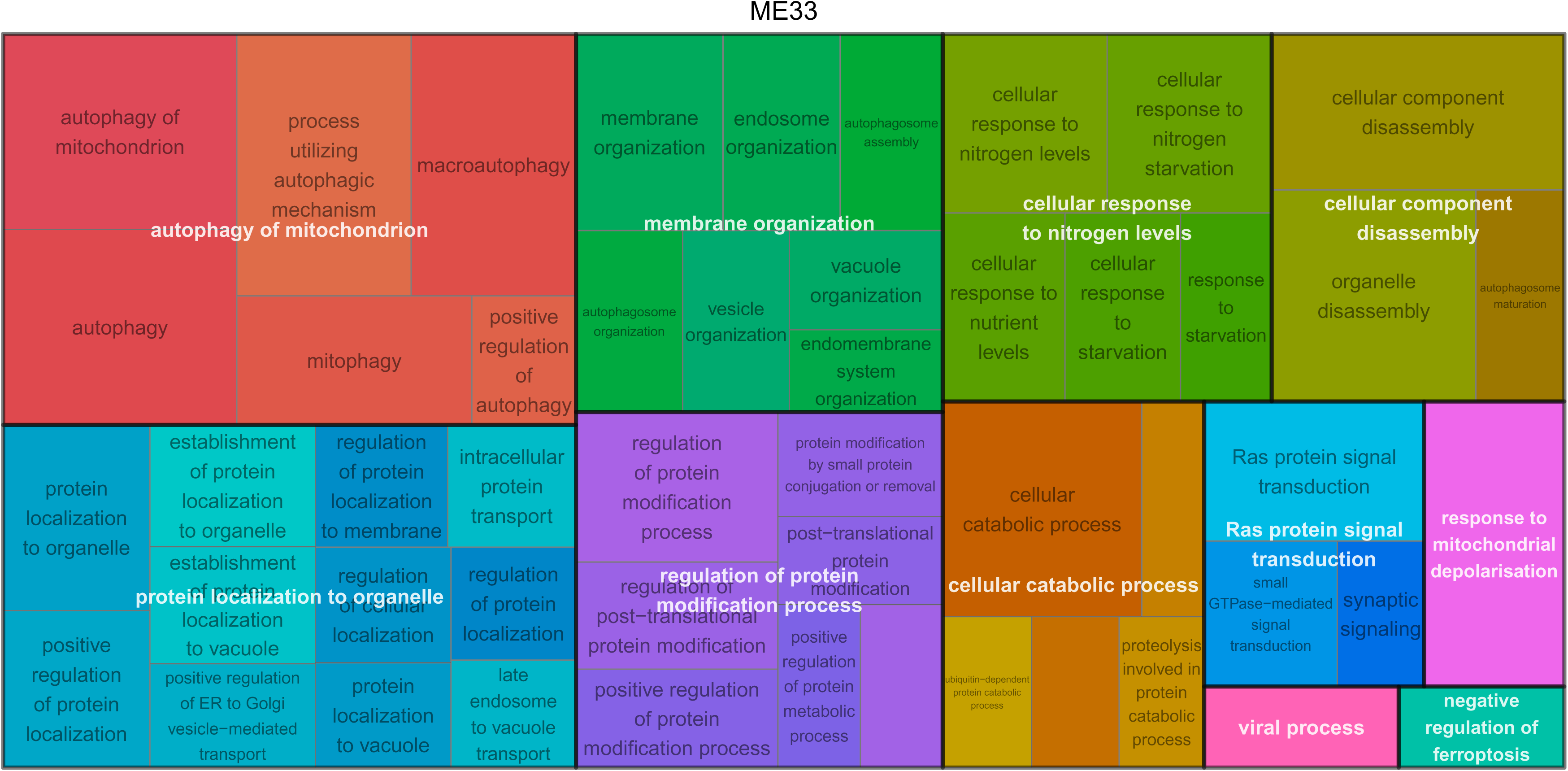

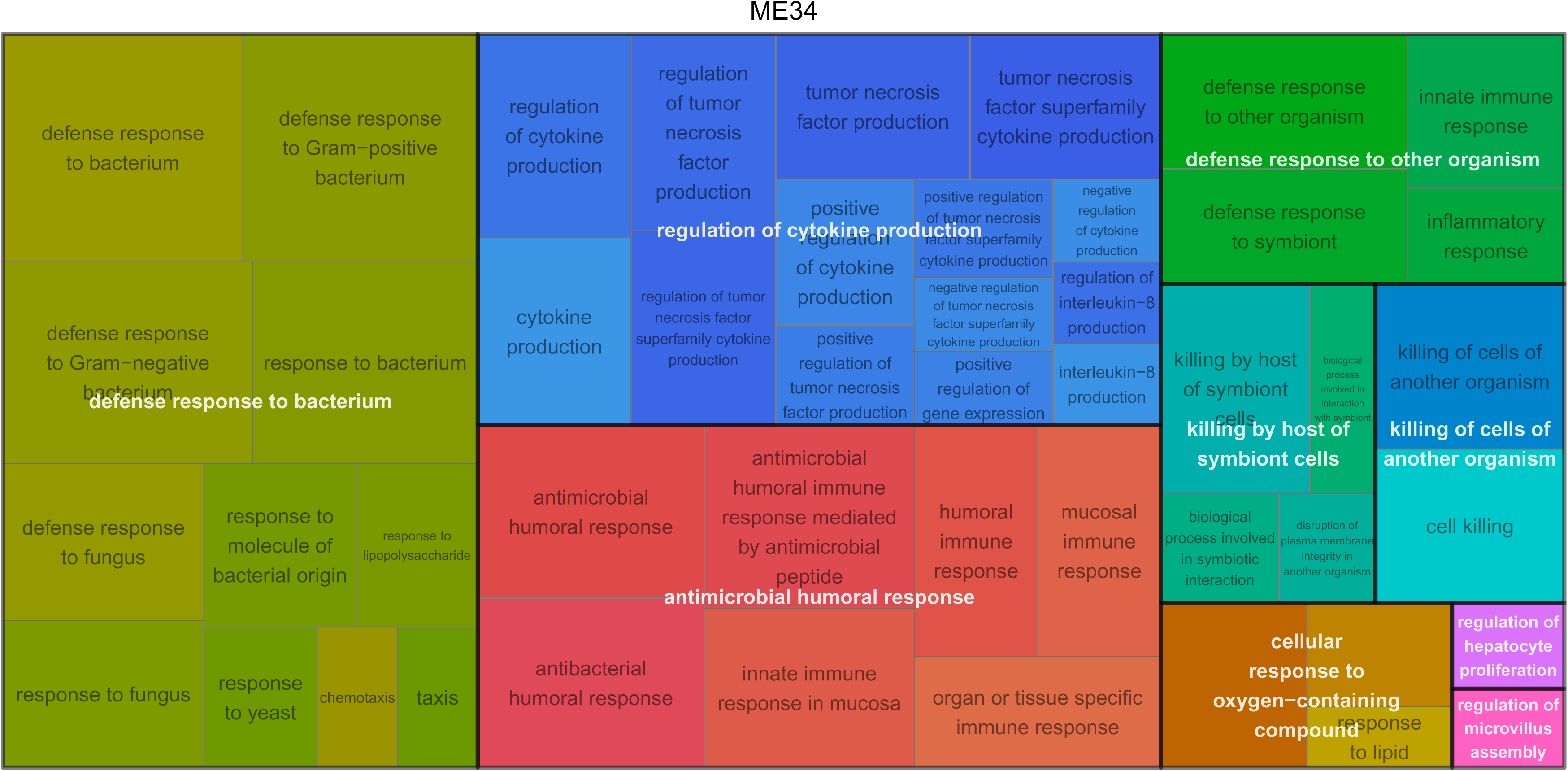

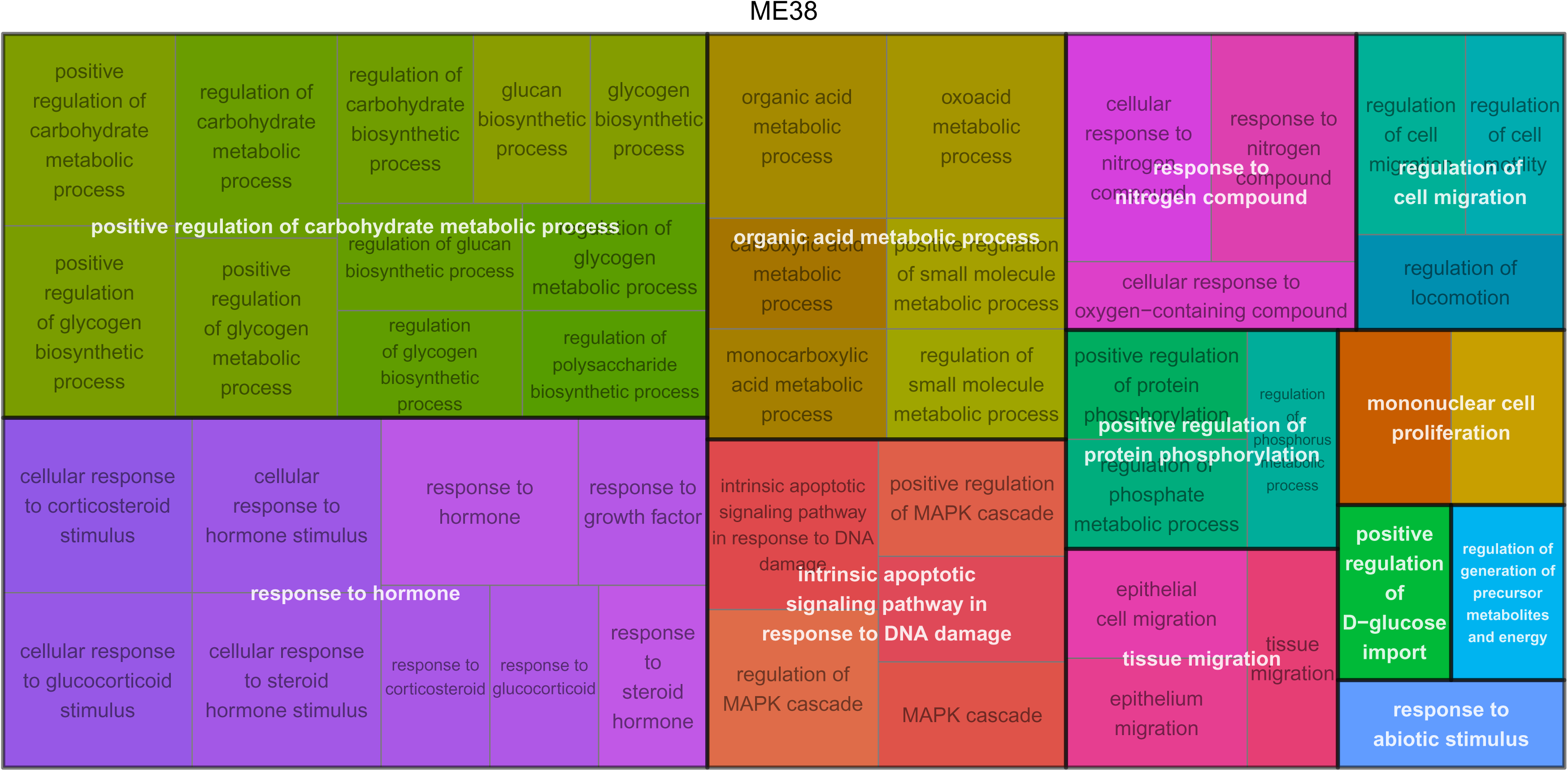

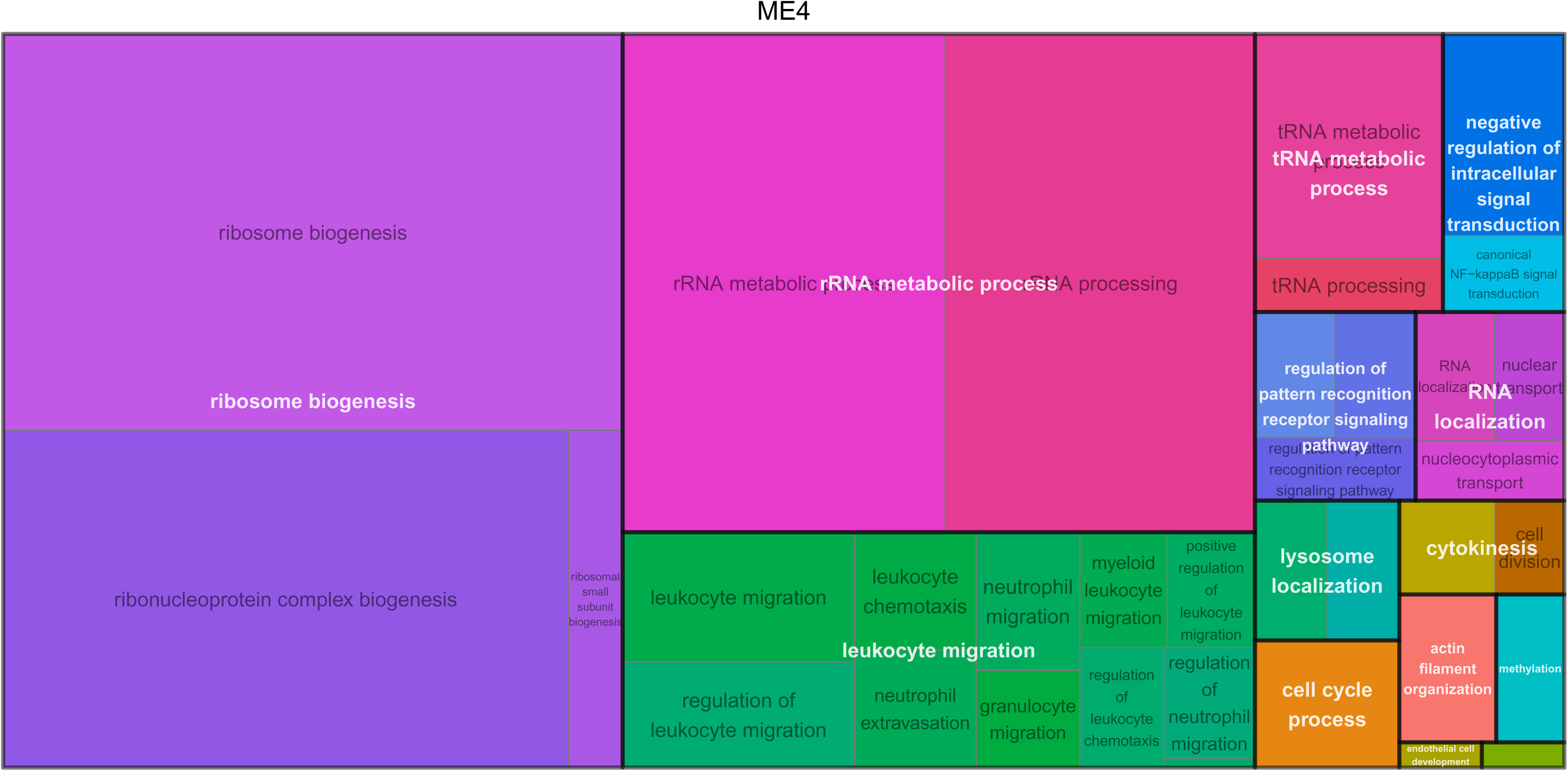

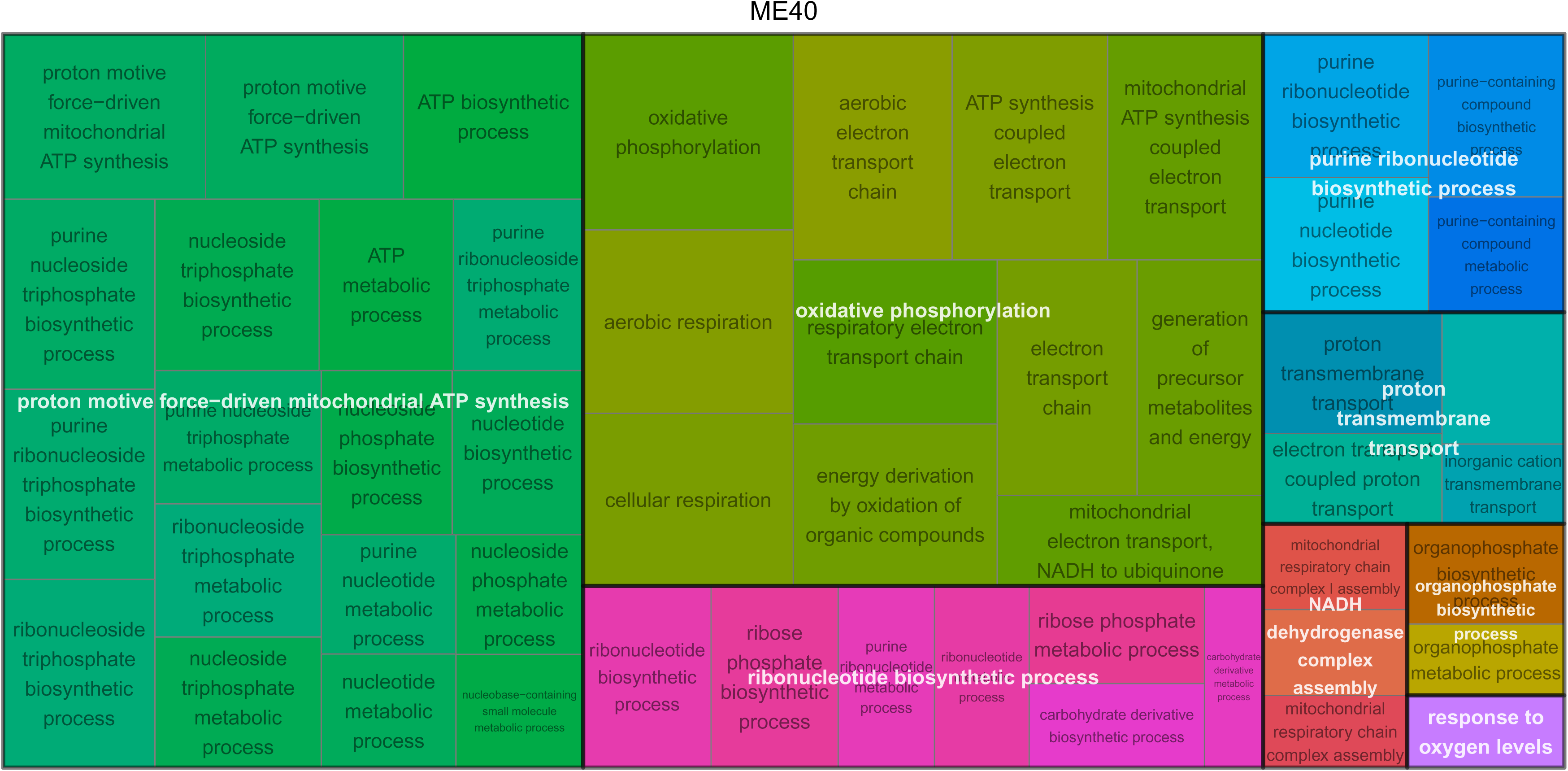

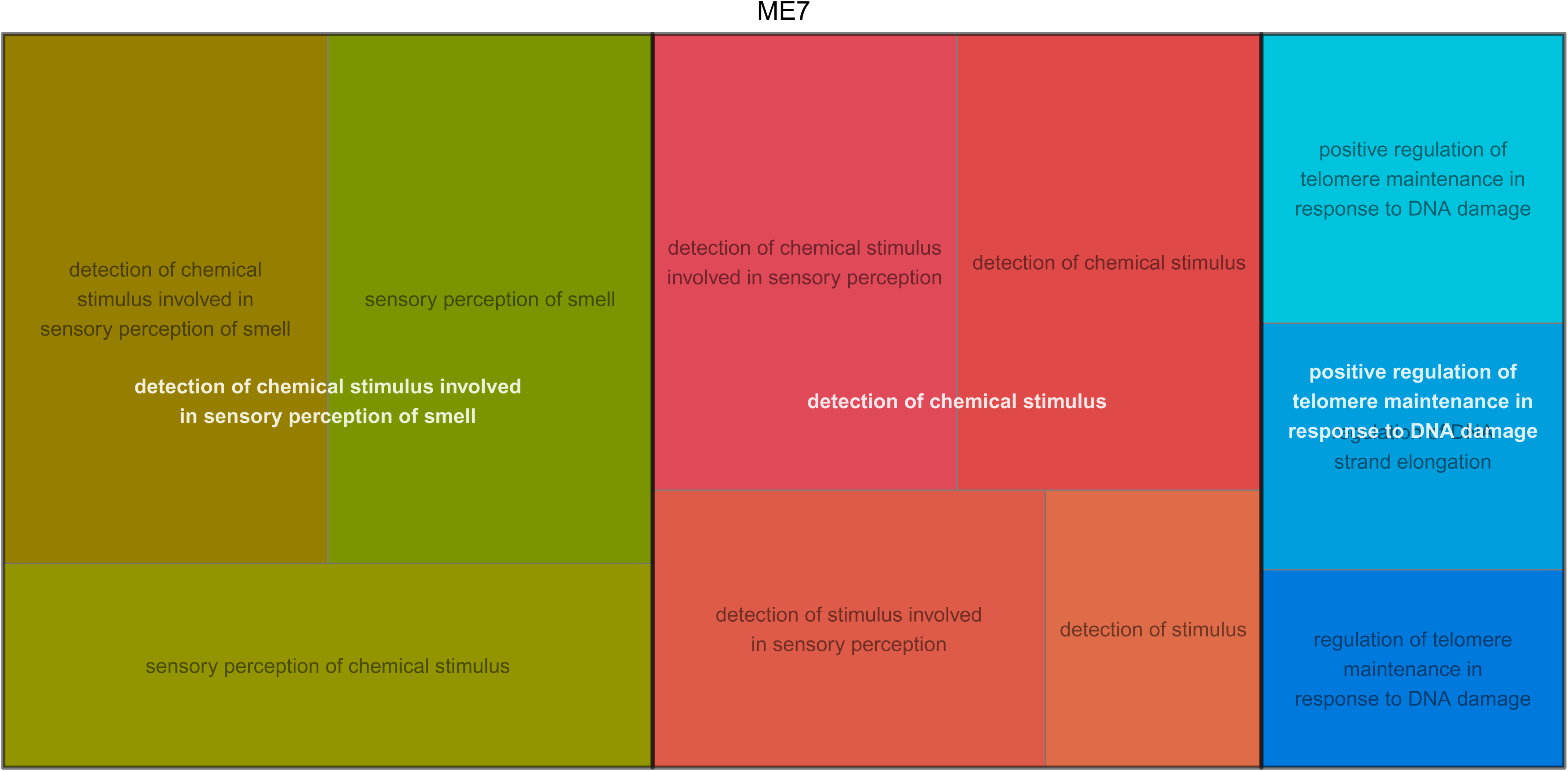

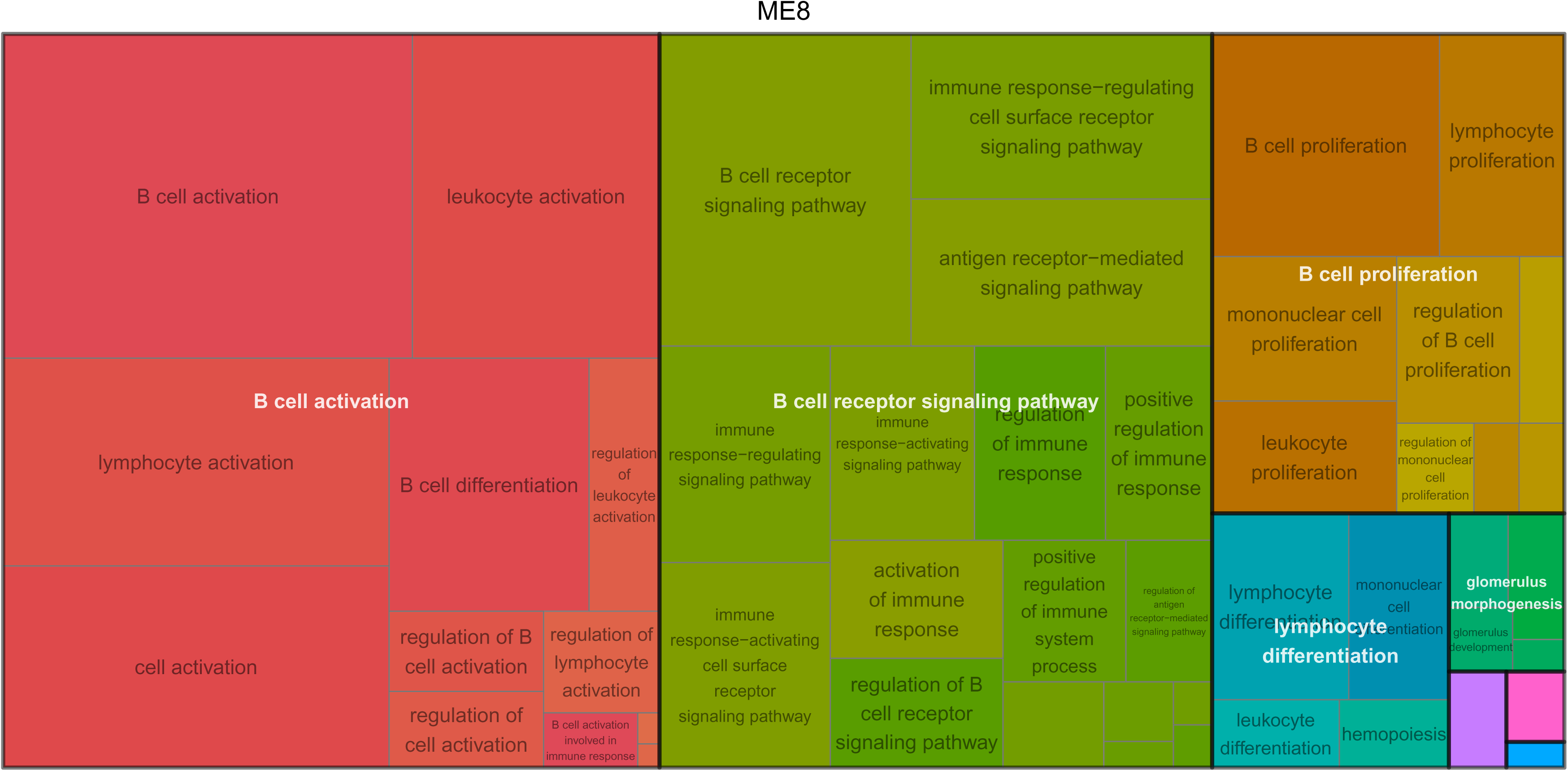

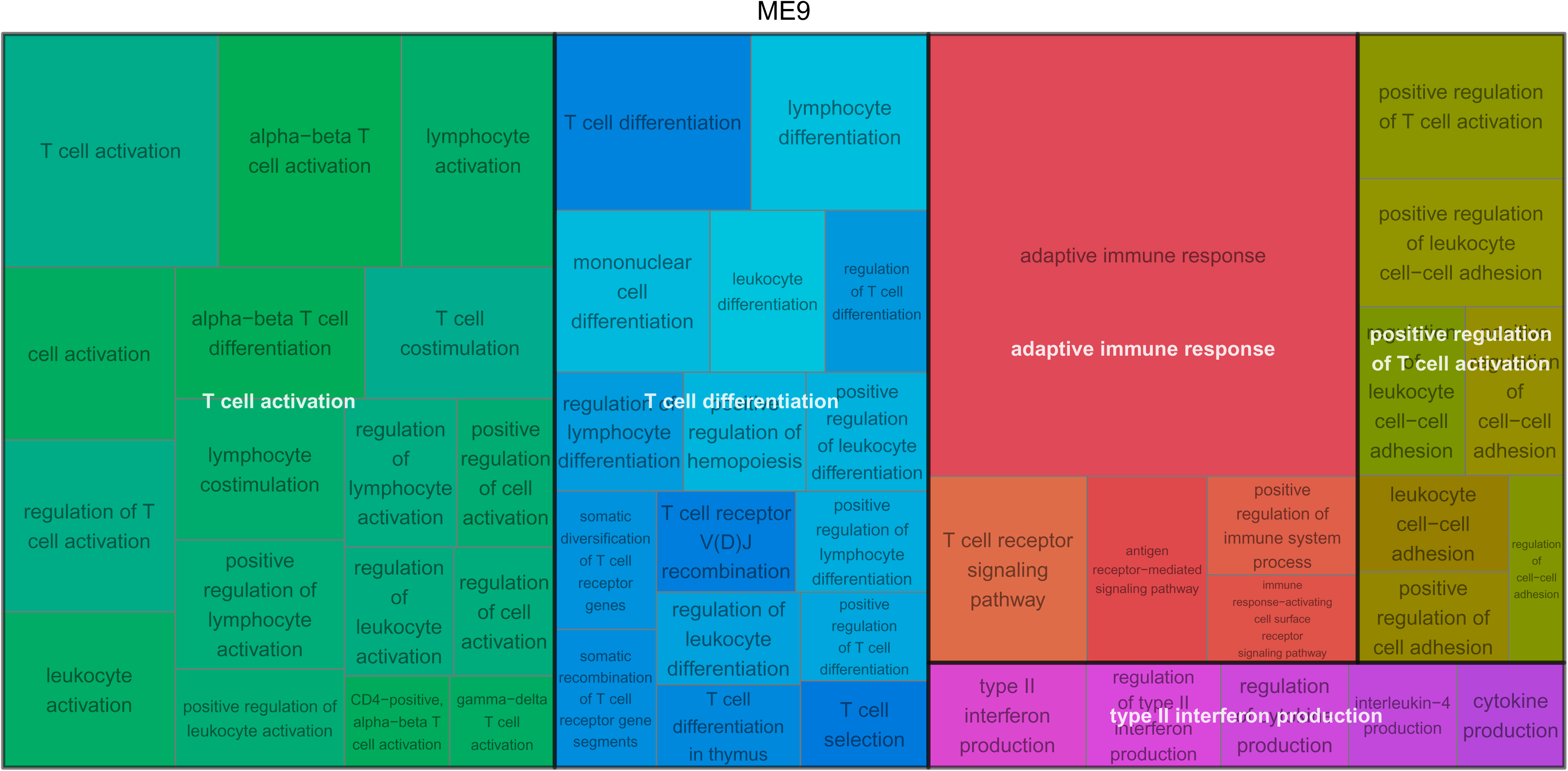

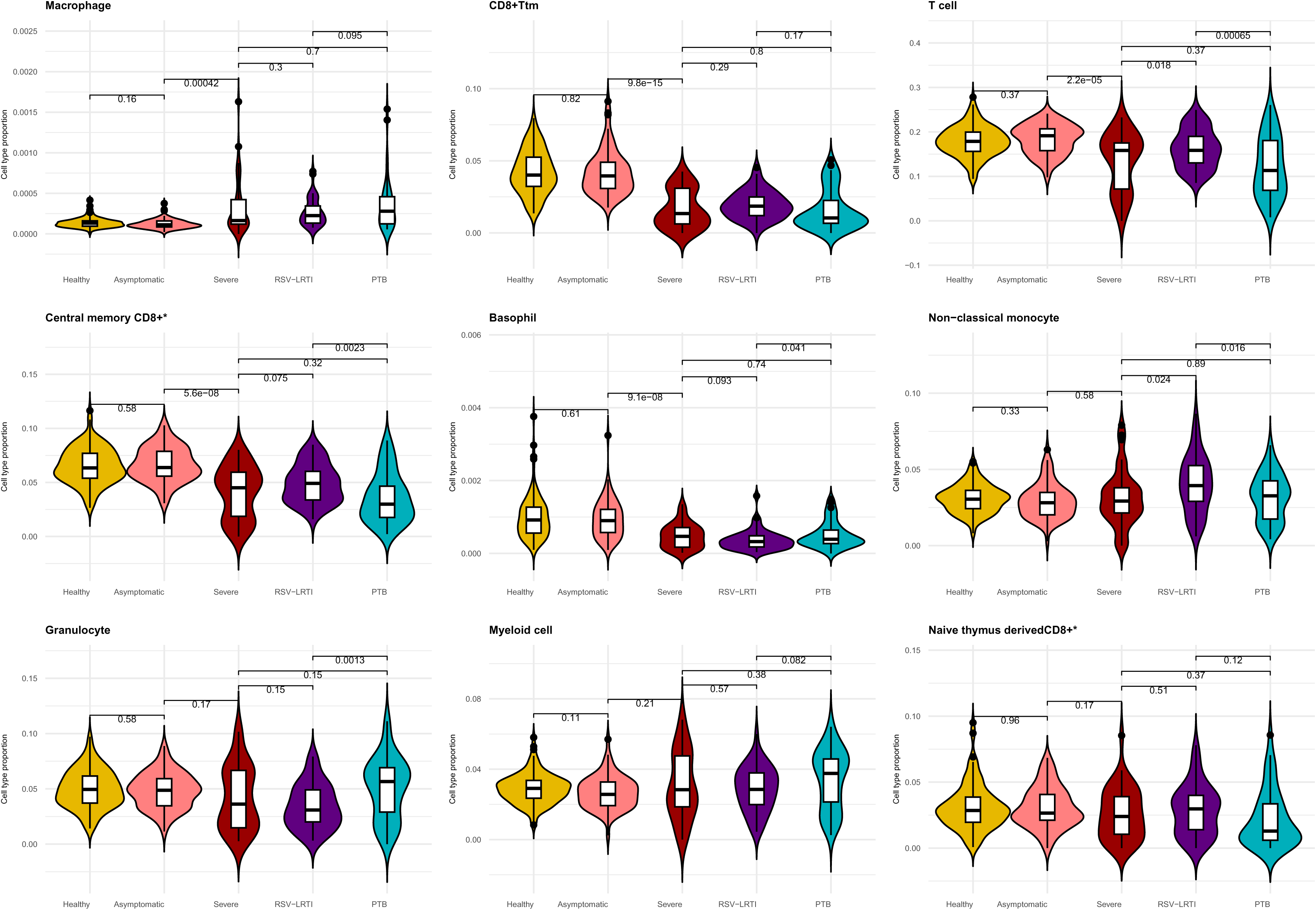

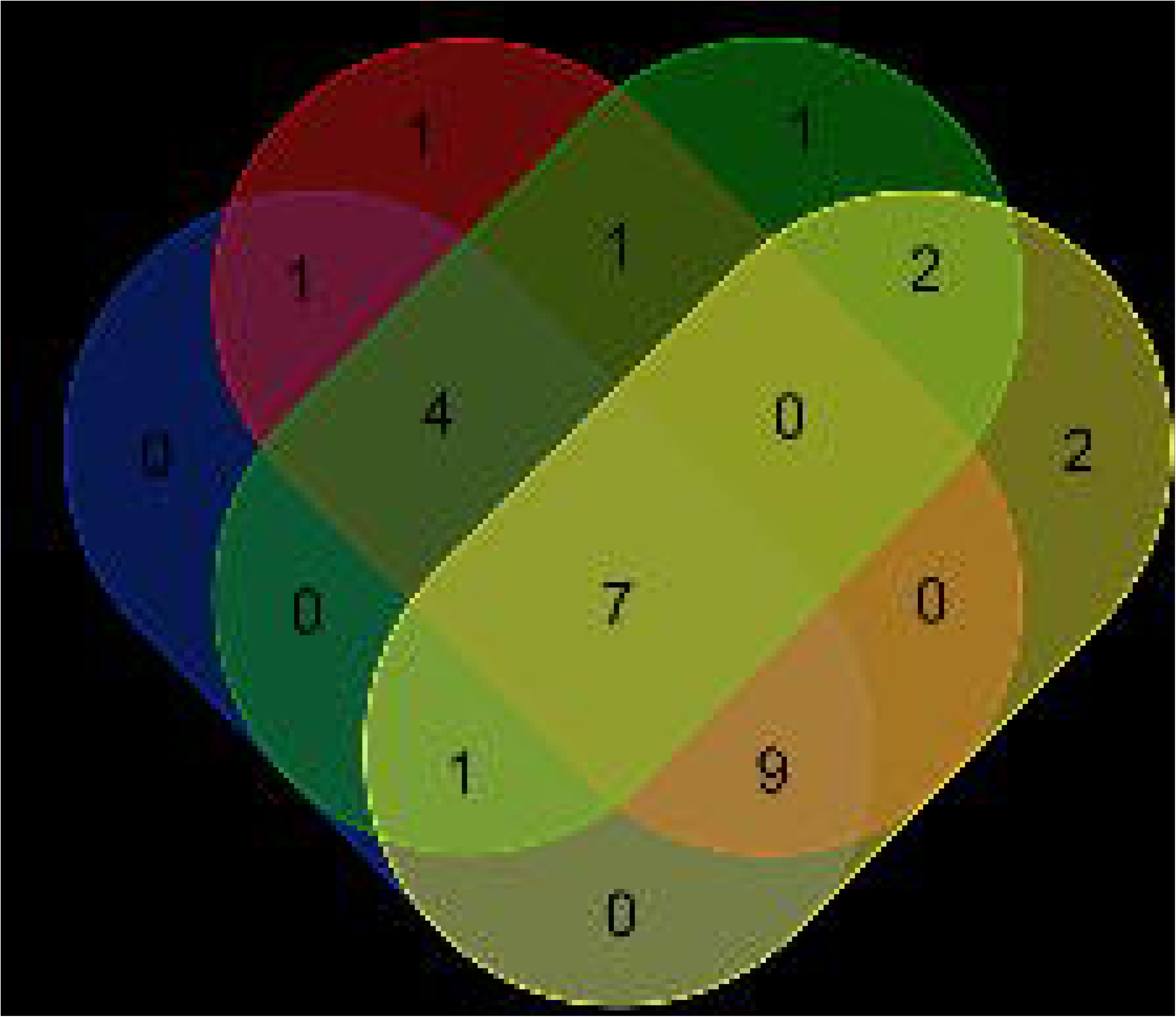

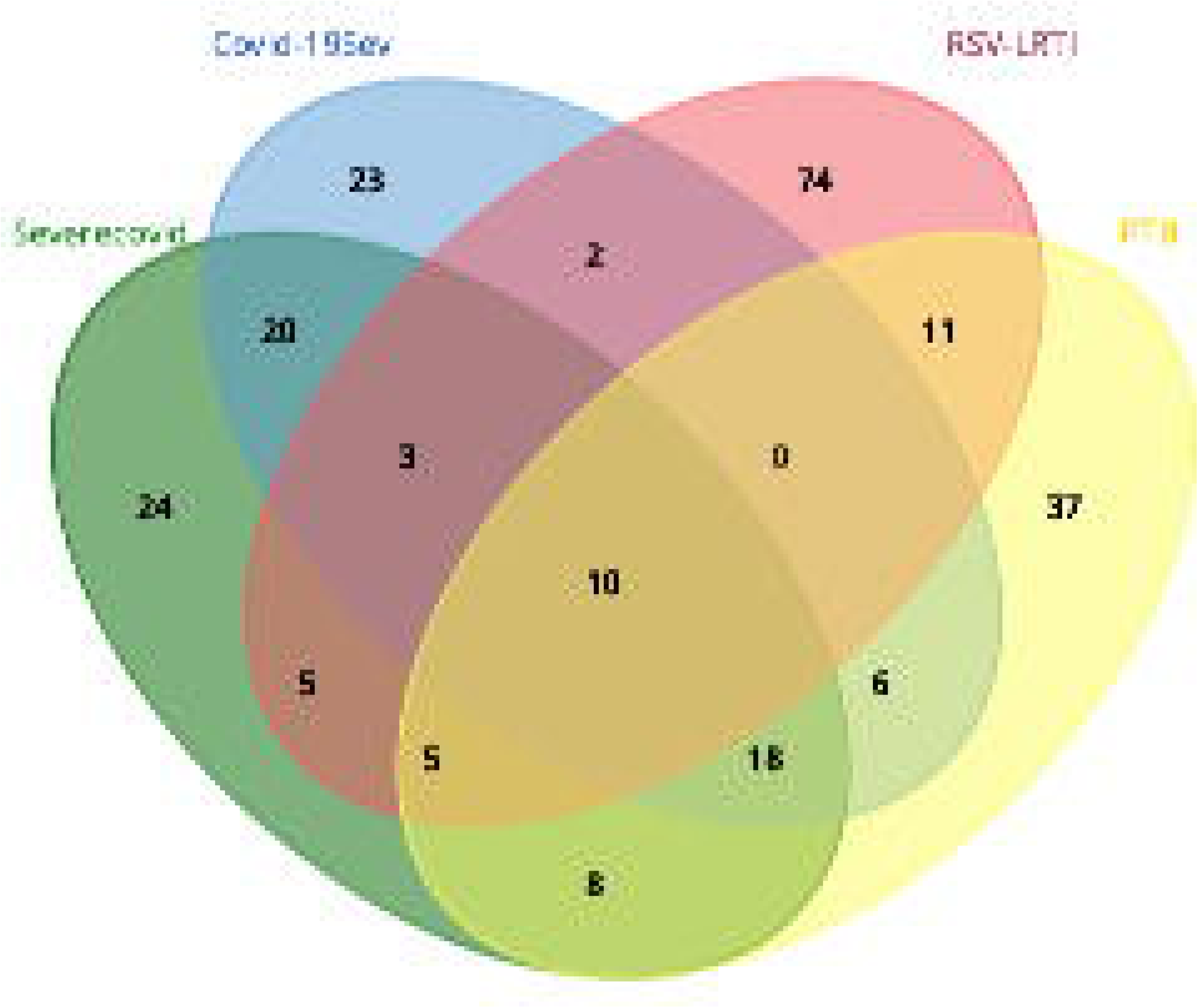

